# Molecular mechanisms of sexual dimorphism in cancer through improved miRNA regulation-level network-based approach: MIRROR2

**DOI:** 10.1101/2025.01.08.631856

**Authors:** Caterina Alfano, Marco Filetti, Lorenzo Farina, Manuela Petti

**Affiliations:** Department of Experimental Medicine, Sapienza University of Rome, Viale Regina Elena, 324, 00161 Rome, Italy; Phase 1 Unit, Fondazione Policlinico Universitario A. Gemelli IRCCS, Rome, Italy; Department of Computer, Control and Management Engineering, Sapienza University of Rome, Via Ariosto, 25, 00185 Rome, Italy

**Keywords:** MIRROR algorithm, differential co-expression network, transcriptomic data, network oncology, sexual dimorphism

## Abstract

A current open challenge in precision medicine is sex-specific medicine: the study of how sex-based biological differences influence people’s health. With recent advancements in high-throughput technologies, large-scale molecular data are being generated for individual cancer patients, however, extracting meaningful insights from these complex datasets remains a challenge. Network-based approaches, being inherently holistic, can lead to a better understanding of the molecular mechanisms underlying a disease. For this reason, this study focuses on the development of a network-based method to investigate sexual dimorphism in cancer using transcriptomic data. Previous studies have already shown how miRNAs are involved in differentiating patients by sex in different types of cancer; however, they focused only on evaluating changes in the expression level, without conducting a more comprehensive analysis of miRNA expression or investigating miRNAs’ targets. The aim of this study is therefore to carry out a multi-layer study involving both miRNAs and their target genes’ expression data. In particular, we developed a generalizable algorithm (MIRROR2), which can be used on cancer patients to help identify key regulatory mechanisms and molecules that act as differentiators between males and females. Here we implemented and tested MIRROR2 on three different cancers (colon adenocarcinoma, hepatocellular carcinoma, and low-grade gliomas) and assessed its performance by comparing it to state-of-the-art approaches. This revealed MIRROR2’s efficacy in identifying sex-specific key genes (and how to integrate them with clinical features), presenting it as a viable alternative to state-of-the-art methods which fail to capture these differences.

## Introduction

Sex and gender specific medicine (SGM) represents a groundbreaking medical approach that asserts the influence of biological sex disparities, gender identity, roles, and relationships on health and illness. These disparities are believed to affect prevention, screening, diagnosis, and treatment; therefore, by learning from and addressing these variations it is possible to enhance healthcare for both males and females^1^. Given that the term “sex” refers to biological/physical differences in the male and female anatomies while “gender” is defined as the socially constructed roles, behaviors, activities, and attributes that a given society considers appropriate for men and women^2^, we can identify two kinds of SGM differences: the sex aspect due to the differences in hormones or genes, and the gender aspect due to the difference in social and cultural roles between men and women^3^. As opposed to sex, which is generally classified as binary, gender is a continuum spectrum; in this study we will only focus on the biological aspects that characterize the male–female dichotomy.

It has been shown that most tumors originating from non-reproductive tissues present biological sex differences, which extend to many aspects of cancer biology and treatment^4^. As a consequence, understanding the leading causes of these disparities is of the utmost importance. In fact, “the interplay between sex chromosomes and hormones influences both local determinants of carcinogenesis, such as cancer initiating cells and components of the tumor microenvironment, and systemic ones, such as cell metabolism and the immune system”. However, limited data exist on how these differences arise at molecular level^1^. By focusing on such a problem, we could identify sex-specific biomarkers for both prognosis and therapy response.

For instance, colorectal cancer (CRC), the third most frequently diagnosed cancer in the world, demonstrates a higher incidence in males compared to females; however, women were also found to have a higher frequency of KRAS mutations in codon 12, which are associated with more advanced adenomas. As of today, the influence of sex on the entire trajectory of colon cancer, from presentation to survival, remains not fully elucidated^5^. As a matter of fact, sex differences in colon cancer have been often attributed to sex hormones, but the molecular mechanisms behind them have not yet been established, and clinical studies are contradictory^6^. Similarly, hepatocellular carcinoma (HCC), which ranks third in mortality rates worldwide, shows a significantly lower incidence in women, suggesting protective effects of estrogens and deleterious effects of androgens. However, the molecular mechanisms of this phenomenon remain to be determined^7^.

The effect of sex has also been studied in the contest of the immune system, proving that this variable affects both innate and adaptive immune responses^8^; however, only a small percentage of immunology-related publications take into account the sex of their subjects, and in immunotherapy clinical trials, women are still underrepresented compared with men^9^. Sex differences in immunobiology are also starting to gain attention in the unique immune microenvironment of the central nervous system (CNS): for tumors like glioblastoma (GBM), where not only there is a gender bias (e.g. secondary GBM, a lower grade glial neoplasm, has a greater incidence rate in women^10^), but almost all patients show recurrence after receiving standard treatment, immunotherapeutic strategies are emerging as a new promising treatment option^11^. Exploring the complex role played by sexual dimorphism becomes a crucial step for the improvement of the therapy response rates.

Many approaches have been explored to study the role of sex in the tumor microenvironment^12–15^, with a recent emerging interest in microRNAs (miRNAs). miRNAs are single-stranded, noncoding RNA molecules (with a length that ranges around 20–24 nucleotides) that play an important role in post-transcription control of gene expression. Specifically, they can induce mRNA degradation or repress translation by binding to the 3’ untranslated region of the target mRNA^16^. "Since the discovery of the first miRNA in 1993 the importance of miRNAs in the regulation of various biological functions including stem cell development and differentiation, organogenesis, early embryo development, and immune responses has been highly appreciated"^17^. The pattern of miRNA expression has also been correlated with cancer type and stage, leading miRNA profiling to be considered an interesting tool for cancer diagnosis and prognosis. Finally, miRNA expression analyses also suggested oncogenic (or tumor suppressive) roles of miRNAs^18^.

In the contest of CRC, for example, knowing that autophagy has a crucial role in cell mechanisms of life and death, it has been possible to identify some miRNAs with an inhibiting effect on autophagy and, as a consequence, a negative influence on the immune response to neoplastic cells^19^. Hasakova et al.^19^, have also seen that the expression of miRNAs is different in males and females; in fact, men show a higher concentration of miR-16, which is associated with a worse survival rate while being downregulated in CRC tissues. Similarly, Lopes-Ramos et al. found substantial differences in gene expression and gene regulatory networks associated with men and women in colon and many other tissues^20^. On the other hand, studies have shown that testosterone induces an upregulation of miR-371a-5p leading to proliferation, neoangiogenesis and metastasis of HCC, which is especially common in men^21^. In women with HCC, instead, it has been identified as an overexpression of miR-18a, with a crucial role for p53: the pathway p53/miR-18a/ER acts just in cells of hepatocellular carcinoma and not in precancerous conditions^22^. Moreover, therapy with estrogens in hepatocellular carcinoma determines an upregulation of the oncosuppressors miR-26a, miR-92, and miR-122 and, consequently, apoptosis. Experiments on rats also found 65 miRNAs to be differentially expressed among men and women; their sex-biased expression was linked with sexually dimorphic molecular functions and toxicological functions that could reflect into hepatic physiology and disease^23^.

This has been the case also for GBM patients, where hsa-let-7c-5p, a crucial miRNA in GBM tumoral mechanism, has been found to regulate cyclin D1 expression through the Wnt/*β*-catenin pathway in osteoblasts; in other words, hsa-let-7c-5p, is a suppressor miRNA with functions in the cancer cell cycle. In addition, Hong et al. have suggested miR-644a as a candidate tumor cell-intrinsic regulator of sex-biased gene expression in GBM as a result of their analysis^24^. Finally, Velázquez-Vázquez et al.^25^ studied the effect of progesterone (P4) on miRNA expression in GBM cells and found that “the possible mRNA targets of the miRNAs regulated by P4 could participate in the regulation of proliferation, cell cycle progression and cell migration of GBMs”.

A limitation of most of the current available studies, however, is to only focus on miRNAs expression, without considering the broader view offered by analyzing miRNAs co-expression patterns and without translating the obtained results into insights that can be used to further continue the analysis at lower-regulatory levels.

In the past, we proposed MIRROR (MIRna RegulatiOn-level diffeRential networks), a new method to investigate miRNAs coordination patterns and how changes in their relationships have an effect at the targets genes’ level that goes beyond quantitative investigations of individual molecules^26^. MIRROR algorithm relies on network theory to analyze the miRNAs differential co-expression given two different subgroups of patient; in particular, it provides a description of the molecular mechanisms that can help explain cancer disparities by projecting miRNA differential networks into the subsequent level of miRNA targets.

In this study we propose an updated version of MIRROR (MIRROR2), which integrates methodological advancements with several novel steps to identify key regulatory mechanisms and molecules that act as differentiators between males and females. Here we provide a detailed description of the MIRROR2 algorithm with methodological and clinical validations, and we show its application to study the sexual dimorphism in colorectal carcinoma, hepatocellular carcinoma and low-grade gliomas, while also comparing it to state-of-the-art methodology.

## Materials and Methods

### Datasets

miRNA and mRNA expression data, as long with clinical and mutational data (simple nucleotide variation) were gathered from The Cancer Genome Atlas (TCGA), through the Genomic Data Commons (GDC) Data Portal R package, by selecting the TCGA-COAD project (colon adenocarcinoma), the TCGA-LIHC project (liver hepatocellular carcinoma) and TCGA-LGG (low grade gliomas). However, mRNA expression of control samples was not available in the TCGA-LGG project; we therefore downloaded the “brain cortex” (Release V8) expression dataset of deceased adult healthy donors from the GTEX Portal.

Clinical features of the patients involved in the study were also retrieved.

For the datasets completely downloaded from the TCGA (i.e., TCGA-COAD and TCGA-LICH), we considered only the patients for which the sex was known and for which miRNA expression of cancer samples and mRNA expression of both cancer and normal samples was available. For the LGG data, we kept all patients for which miRNA and mRNA expression of cancer samples was available and we added to them the mRNA control cases downloaded from the GTEX portal.

Mature miRNAs accession numbers were translated into miRBase IDs with the miRBaseConverter R package^27^.

For both mRNA and miRNAs, we filtered out low-count values by keeping only the ones with a count greater or equal to 10 in at least 90% of the available samples. We then applied Deseq2^28^ normalization on the raw counts.

## Methods

### Co-expression networks

A co-expression network is a graph *G*(*M*, *E*) consisting of a set of vertices (or nodes) *M* and a set of edges *E*. Each *m* ∈ *M* represents a transcript (miRNA or mRNA), whereas each *e* ∈ *E* represents a significant co-expression relationship between two transcripts. The co-expression across samples is typically measured with a correlation coefficient such as Spearman or Pearson rho.

### Differential co-expression network

A differential co-expression network is a graph *F*(*M*, *Z*), where *M* is the same set of nodes of two co-expression networks associated to two groups of samples (*N*_1_, *N*_2_) to be compared, while *Z* is a set of edges in which each *z* ∈ *Z* represents the statistical change in the co-expression of two transcripts between the datasets under investigation. The differential network can be obtained with the diffCorr^29^ methodology; this consists in applying the Fisher-Z transformation to the correlation coefficients (*E*1, *E*2) and computing z-scores with the following formula:

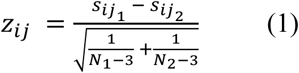

where *S*_*ij*_1__ and *S*_*ij*_2__ are the Fisher-Z transformation of, respectively, the correlation in the co-expression network *G_1_* and in the co-expression network *G_2_*, while *N_1_* and *N_2_* are the number of samples in each group of samples. Positive z-scores represent increases of correlation in the first network, while negative z-scores represent increases of correlation in the second network. A thresholding step is typically applied to consider only significant changes in co-expression. The resulting network is undirected, weighted and signed, with the link’s sign coding for the condition in which the correlation is greater. A positive edge weight means that the co-expression between the two nodes is stronger in the first condition and vice versa.

### MIRROR2 algorithm

In this study we propose a new method to investigate MIRna RegulatiOn-level diffeRential networks (MIRROR2) which is based on the computation and analysis of co-expression and differential co-expression networks. The proposed algorithm comprises two main phases (Fig. 1):

1. miRNA differential network analysis to compare co-expression patterns in males and females
2. sex-specific mRNA differential network analysis to compare healthy and cancer samples based on the subset of miRNA identified in phase 1.

**Figure 1.**
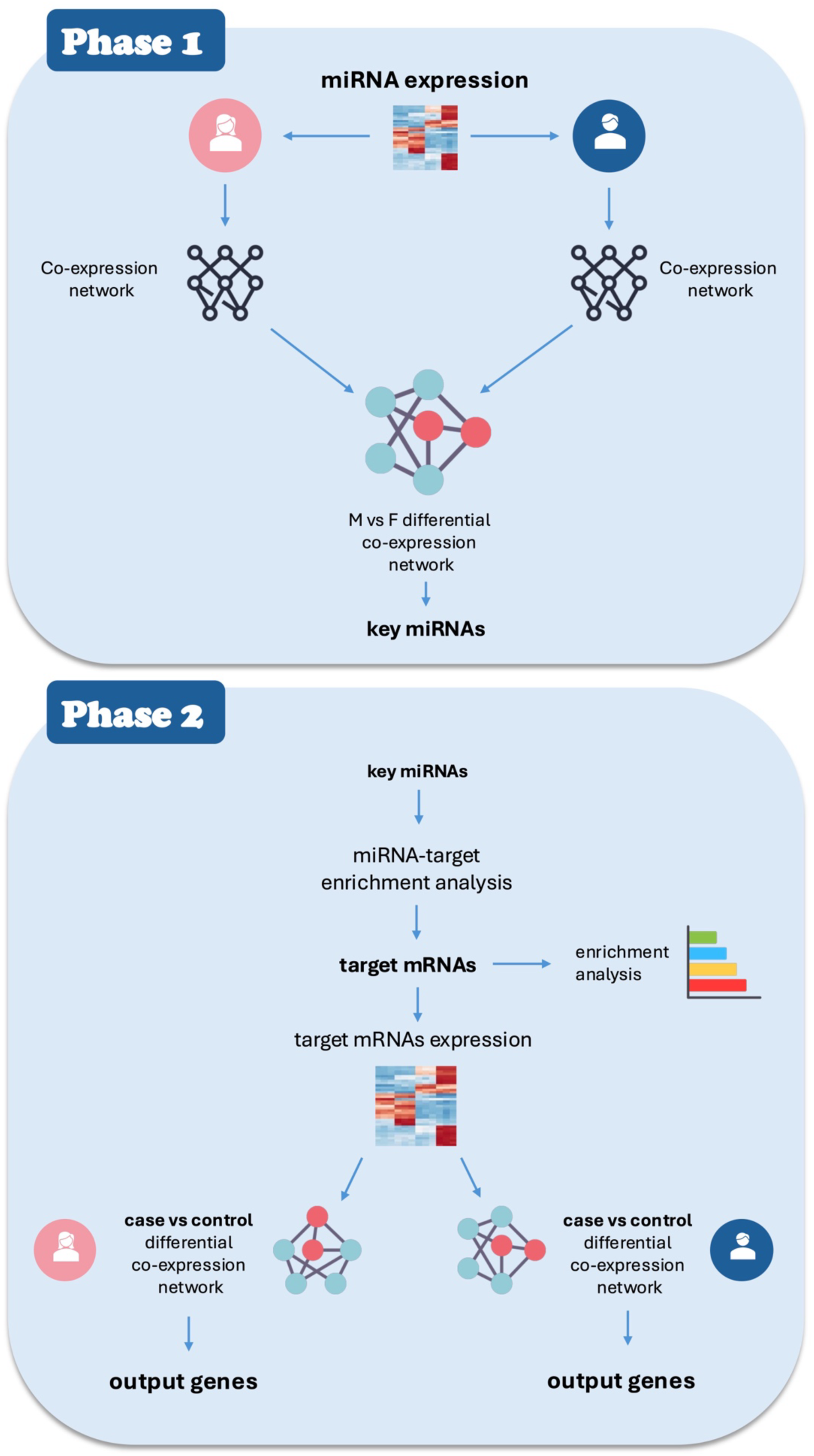
Overview of MIRROR2’s workflow

#### Phase 1

The inputs of the first phase of MIRROR2 are two miRNA expression datasets (one belonging to male patients and one belonging to female patients) with size *m x N_k_*, where *m* is the number of miRNAs and *N_k_* (with *k=1,2*) is the number of patients of each group. Sex-specific co-expression networks (*G_M_* and *G_F_*) are then computed by assessing the correlation of the available miRNAs within each cohort of patients. Then, comparing the two co-expression graphs by assessing the difference in correlation, the differential co-expression network *F_MvsF_* is obtained. MIRROR2 identifies the key players that drive the differences in males and females at regulatory level by analyzing the differential network. Steps included in the topological analysis of this network are: the computation of network density, the extraction of the largest connected component and the identification of the most important nodes. In this application, we consider degree centrality as the most important characteristic of a node: degree counts how many neighbors a node has and thus, given a miRNA, informs how many other miRNAs it changes its co-expression relationship with based on sex.

The output of Phase1 is a set of miRNAs with a crucial role in differentiating males and females in terms of co-expression patterns. Indeed, MIRROR2 identifies these miRNAs by selecting the most connected nodes of the differential network (95th percentile of the degree distribution) and then expands this set by adding their first neighbors.

#### Phase 2

The first step of the second phase consists in finding the target genes associated with the set of key miRNAs received as input. For this purpose, since a single miRNA may have hundreds of different targets and a given target might be regulated by multiple miRNAs, MIRROR2 performs a miRNA-target enrichment analysis (hypergeometric test with FDR correction, adjusted p-values < 0.01) to prioritize miRNA-target interactions found in the mirTarBase database. In this way we obtain the set of genes *T* on which the selected miRNAs can exert a differential regulatory action based on the subtype of patients (male/female). To characterize the target genes obtained and inspect their functions, MIRROR2 then performs an enrichment analysis to highlight the pathways and biological processes in which these genes are involved. Then, similarly to what has been previously done with miRNAs, these target genes are used to obtain mRNA co-expression networks. In this phase, the goal is to compare cancer samples with control samples for male and females separately. For this purpose, MIRROR2 computes a total of four co-expression networks:

- *G_CM_*(*T*, *E*) for cancer samples of male patients
- *G_NM_*(*T*, *E*) for control samples of male patients
- *G_CF_*(*T*, *E*) for cancer samples of female patients
- *G_NF_*(*T*, *E*) for control samples of female patients

By comparing these networks, two additional differential co-expression networks are obtained:

- *F_CvsN_M_*(*T*, *Z*) comparing cancer vs control samples of male patients
- *F_CvsN_F_*.(*T*, *Z*) comparing cancer vs control samples of female patients

In all these graphs, *T* represent the set of target genes.

These differential networks are a mean to highlight how gene co-expression changes between normal and cancer samples; having separate networks for males and females allows to further investigate the sex-biased differences of the tumorigenic process. Node strength (i.e., the sum of the weights of node edges) is then computed. In particular, the absolute values of the z-scores computed to build the network are used as the weights of node edges:

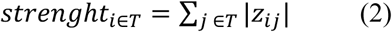

Finally, MIRROR2 selects the nodes (i.e., target genes) with highest strength from the two networks and returns them as output.

#### Performance evaluation

To assess our algorithm’s performance, we performed different validation analyses. At first, we performed a comparative analysis with other two well-established bioinformatics approaches: differential expression analysis and weighted correlation network analysis (WGCNA)^30^. Moreover, to assess the sexually dimorphic specific alterations our miRNA differential co-expression network, we run a permutation analysis to obtain networks made from shuffled data.

These validation steps were performed on all three datasets.

Finally, a clinical validation was carried out on the COAD dataset to evaluate the potential clinical relevance of the output genes of MIRROR2.

#### Comparative analysis with other approaches

Given that one of the main characteristics of MIRROR2 is to start the analysis from miRNA expression data, differential expression and the WGCNA were both applied both to the input miRNAs, and their output compared to MIRROR2 Phase1 output. Differential expression was also evaluated for the mRNAs identified in Phase2.

### Weighted correlation network analysis (WGCNA)

The WGCNA is a widely used bioinformatics software that is applied to analyze high-dimensional gene expression data, comparing samples of different types with the aim of identifying key molecules to explain the examined conditions. This method creates scale-free co-expression networks and identifies genes clusters that can be analyzed to understand the genes’ functions and to discover relationships between genes and phenotypes of interest. The most representative genes of each module, called module eigengenes (MEs), were identified by taking the first principal components. MEs are then used to compute module trait correlation, a measure that allows us to relate how much a module is associated to a phenotype. Finally, module membership (MM) and gene significance (GS) were computed for the genes belonging to the modules of interest. Module membership is measured as Pearson’s correlation coefficient of the expression profile of one gene in all samples and the ME of the module it has been assigned to. GS, instead, represents the Pearson’s correlation coefficient of singular genes’ expression profiles to the trait of interest.

### Differentially expressed genes (DEGs) and miRNAs

To assess differential expression among two sets of patients, for each miRNA /mRNA we computed the fold change i.e., the ratio between the mean expression in the first set and the mean expression in second set in a log2 scale. Then, to assess the significance of the fold change values we performed a student t-test with correction for multiple comparisons. To identify differentially expressed genes and miRNAs, 1.5 was chosen as a threshold for the fold change absolute value, while 0.01 was selected for the significance level of the adjusted p-value.

#### Evaluation of robustness with respect to the sex variable

A computational validation was made to ensure that the network and miRNAs identified with MIRROR2 are specific to the sex-based split of the datasets. To do this we run 1000 simulations in which we shuffled patients and divided them, regardless of the sex variable, into groups with the same size as the original dataset. We then inferred the miRNA differential co-expression network in each simulation and retrieved the most connected miRNAs as well as their first neighbors. We compared the miRNAs obtained in each simulation with the miRNAs obtained by MIRROR2 on the real dataset by computing the Szymkiewicz–Simpson overlap coefficient:

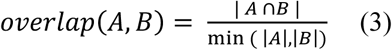

whit *overlab(A,B)* ∈ [0,1]. If set A is a subset of B or the converse, then the overlap coefficient is equal to 1. We then computed the median overlap across all simulations. Moreover, we also assessed the overlap significance by applying the hypergeometric test to see if the miRNAs obtained in the simulation were statistically associated in the original miRNAs. P-values obtained from the hypergeometric test were then corrected with the Bonferroni method and considered significative when lower than 0.01.

#### Clinical validation

A final validation was done to show the potential of using the output genes of MIRROR2 in a clinical setting. As a demonstrative example, we used the output genes of the COAD application to make a binary classifier which can predict if a patient will have a short or long survival time. All the patients for which mRNA expression and survival outcome was available in the TGCA-COAD dataset were retrieved, for a total of 207 females and 234 males; then high or low survival time for each patient was defined by setting a threshold of 24 months, a value close to the median survival time, so to have a balanced amount of the two classes. The 20% of both datasets (male and females) was removed and kept as a test set, while the remaining 80% was used to build the models. Two logistic regression models were trained to create predictions based on all the final output genes: one for females (based on the genes extracted from the female differential co-expression network) and one for males (based on the genes extracted from the male differential co-expression network). We then assessed these models’ performances by computing the ROC curve, the F1 score and the confusion matrix. Finally, to check that the genes are sex specific, we tried to apply the model trained on female data to male patients and vice versa and checked their predictive abilities.

## Results

### Colon adenocarcinoma

The colon adenocarcinoma input dataset is made of 231 miRNAs and 38 patients, equally split into 19 males and 19 females. The two subgroups of patients do not show significant differences in terms of clinical characteristics or survival (Supplementary Figure 1). Mutational data shows cohesive variants and SNV among the two sexes, while the top mutated genes are mostly different (Fig. 2 and Fig. 3).

**Figure 2.**
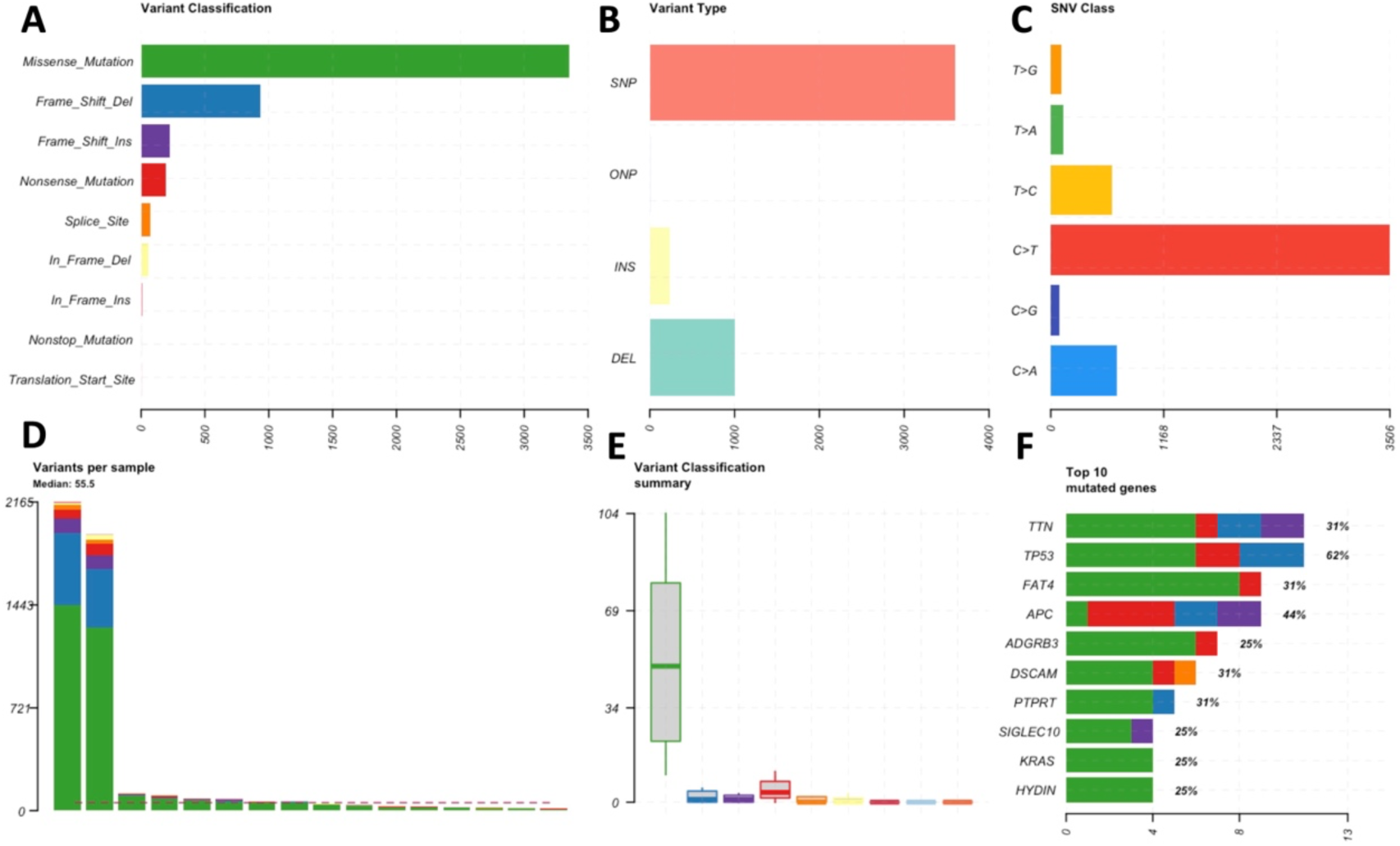
Summary of mutational characteristics of female samples. It includes barplots accounting for number of variants by class (A), type (B) and single nucleotide variation class (C) found in the samples. D) Number of variants per sample (from most to least mutated). E) Boxplots of the number of variants by class, without outliers. F) Top 10 mutated genes, ordered by number of mutations, with percentage of samples in which they are mutated.

**Figure 3.**
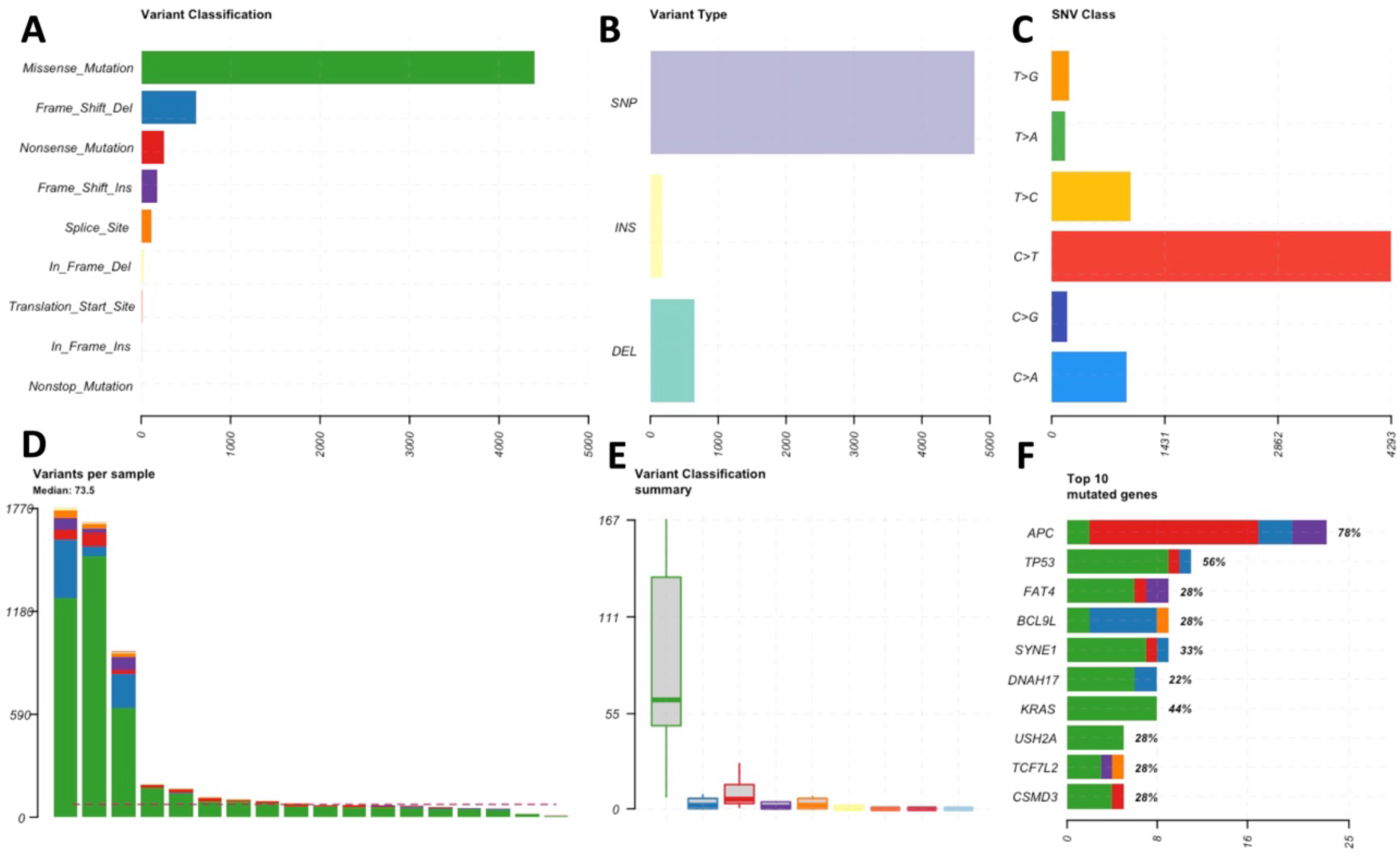
Summary of mutational characteristics of male samples. It includes barplots accounting for number of variants by class (A), type (B) and single nucleotide variation class (C) found in the samples. D) Number of variants per sample (from most to least mutated). E) Boxplots of the number of variants by class, without outliers. F) Top 10 mutated genes, ordered by number of mutations, with percentage of samples in which they are mutated.

#### MIRROR2 phase 1

The miRNA differential network (Fig. 4) has a density of 0.011, where 47% of the edges are negative (representing correlation that are stronger in women) and 53% are positive (correlation that are stronger in men). We then identified 5% most connected miRNAs: miR-101-3p, let-7i-5p, miR-134-5p, miR-29c-3p, miR-339-5p, miR-429, miR-451a, miR-146b-5p, miR-146b-3p. By gathering first neighbors of these 9 miRNAs, we identified 89 miRNAs of interest to serve as input for the second phase of the algorithm.

**Figure 4.**
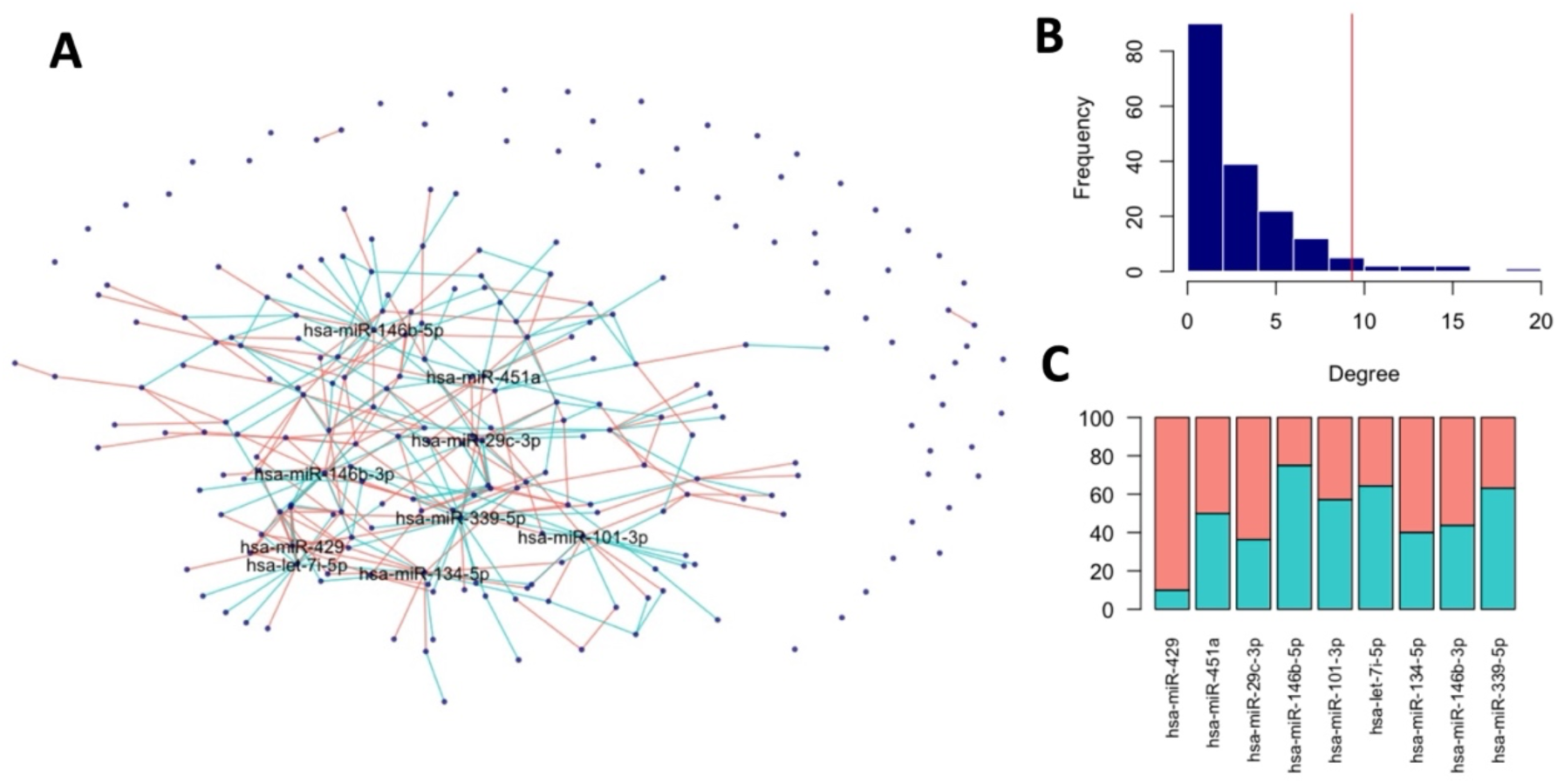
A) miRNA differential co-expression network among males and females. Each node is a miRNA, and each edge is a significant change in co-expression. Blue edges code for a positive differential co-expression (meaning that co-expression is stronger in males) while pink edges code for a negative differential co-expression (meaning that co-expression is stronger in females). B) Degree distribution of network’s nodes. The red vertical line signals the 95th percentile of the distribution. C) Degree composition of the most connected nodes, showed as percentage over total degree.

#### MIRROR2 phase 2

Target enrichment analysis of the 89 miRNAs revealed 257 genes, 241 of which were present in our mRNA expression dataset. These target genes are enriched in KEGG pathways that have an important role in colon cancer and are being studied in the context of sexual dimorphism^31–33^, such as the Estrogen Signaling Pathway (adj p-val = 3.4 · 10^-6^), Proteoglycans in cancer (adj p-val = 8.6 · 10^-11^) and Ras signaling pathway (adj p-val = 8.8 · 10^-7^). When inferring differential co-expression networks of case and control samples for males and females separately, we obtained two networks with different density (0.04 for males and 0.1 for females), and different degree distribution.

As we expected, the sets of genes with higher node strength for male and female samples turned out to have little to no overlap (Fig. 5A). Both networks do, however, have a balanced amount of positive and negative edges.

**Figure 5.**
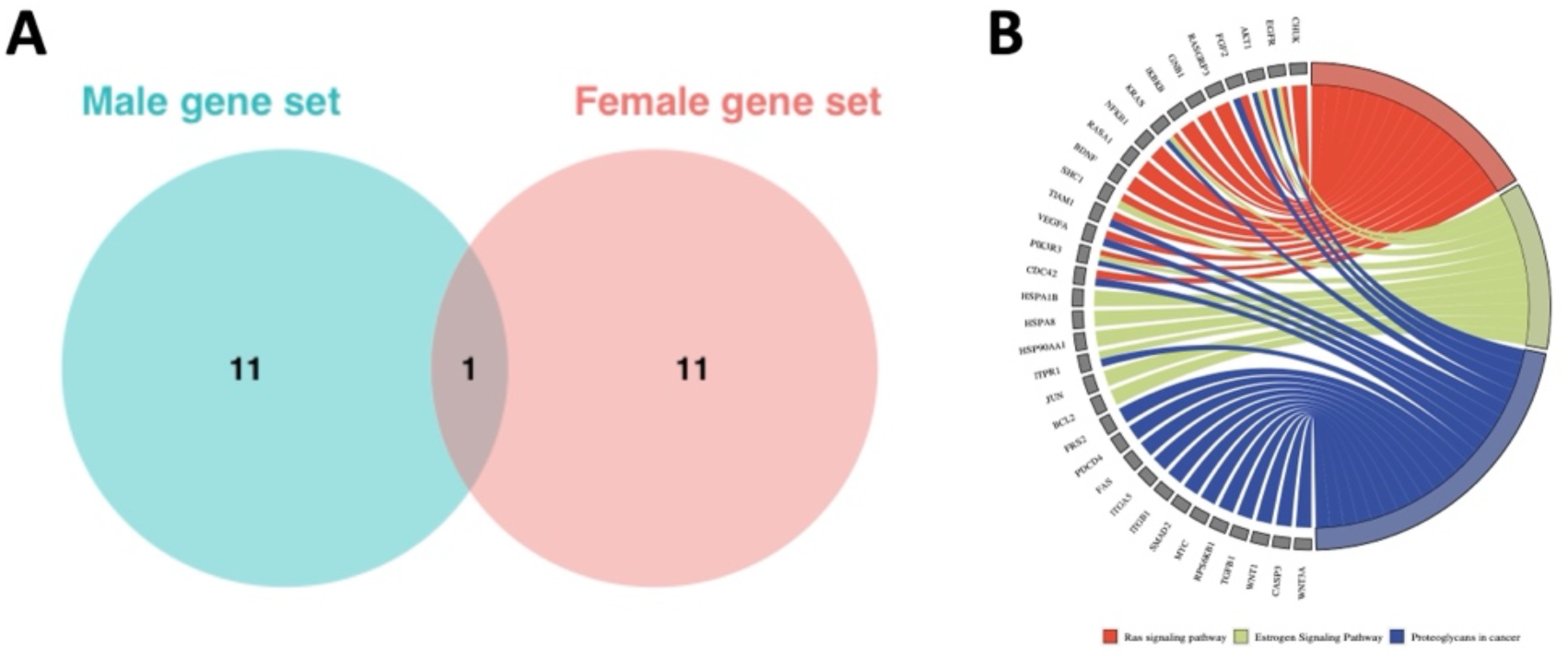
A) Venn diagram showing the overlap between the most connected nodes of the female and male differential co-expression network. B) Chord plot showing the genes belonging to KEGG pathway of interest for this application.

The genes with higher strength in the female differential network are: RAB11FIP2, TUBB4B, CCNE2, AKT1, GNB1, TRAP1, CALU, ANKRD12, USP8, USP48, PSMD2 and NRBP1. Instead, for the male differential network, they are: QKI, XPO1, RAB11FIP2, BTBD7, PTGS2, SMAD2, SNX16, TADA2B, ERMP1, ARHGAP32, YBX3 and NFIA. These two gene sets are the final output of the MIRROR2 algorithm.

### Hepatocellular carcinoma

The hepatocellular adenocarcinoma input dataset is made of 551 miRNAs and 49 patients (27 males and 22 females). Patients do not show significant differences in terms of clinical characteristics or survival (Supplementary Figure 2) based on their sex. A similar mutational profile is shown by analyzing variants and SNV among the two sexes; some of the top mutated genes are also common (Fig. 6 and 7).

**Figure 6.**
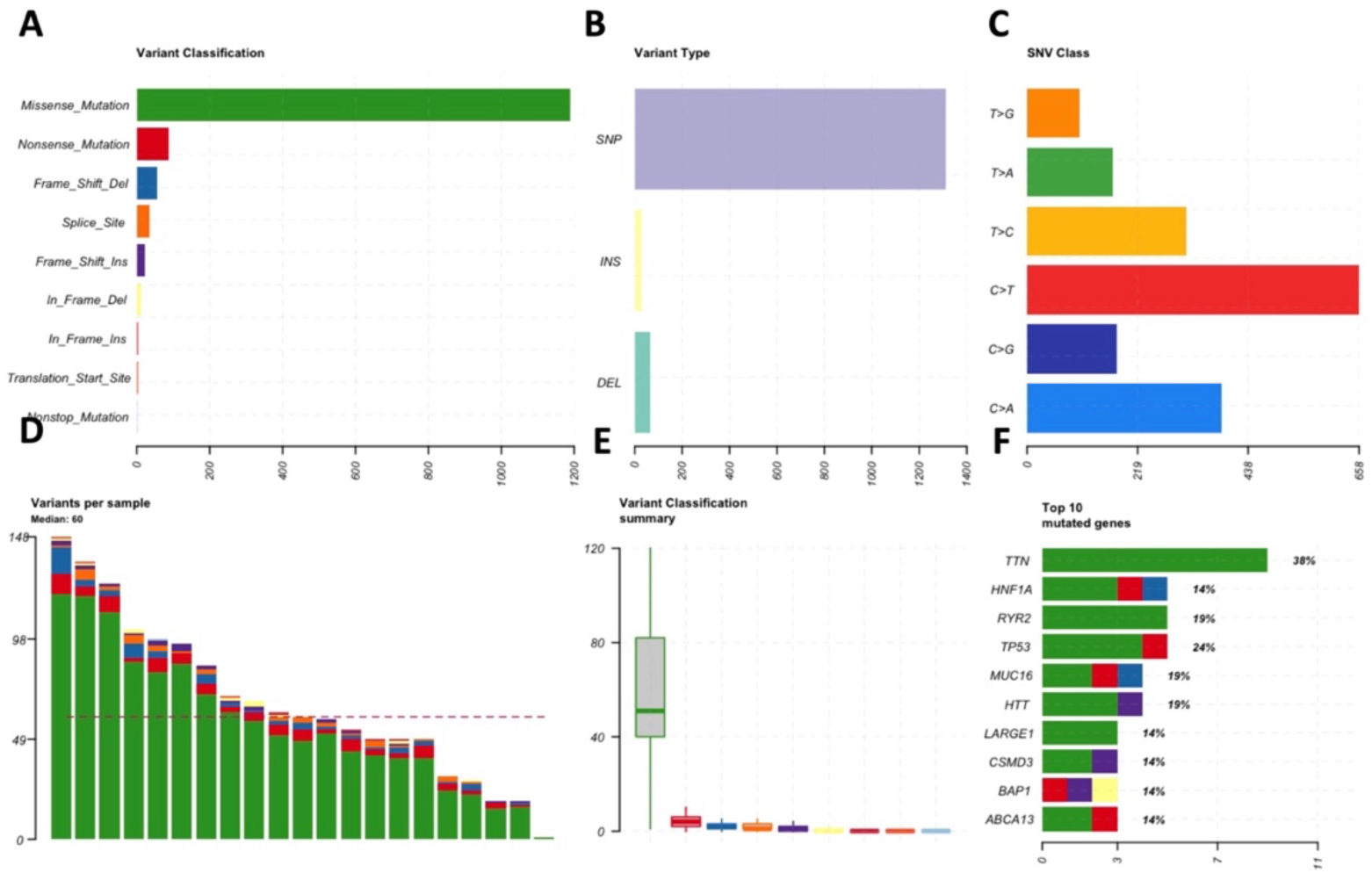
Summary of mutational characteristics of female samples. It includes bar plots accounting for number of variants by class (A), type (B) and single nucleotide variation class (C) found in the samples. D) Number of variants per sample (from most to least mutated). E) Boxplots of the number of variants by class, without outliers. F) Top 10 mutated genes, ordered by number of mutations, with percentage of samples in which they are mutated.

**Figure 7.**
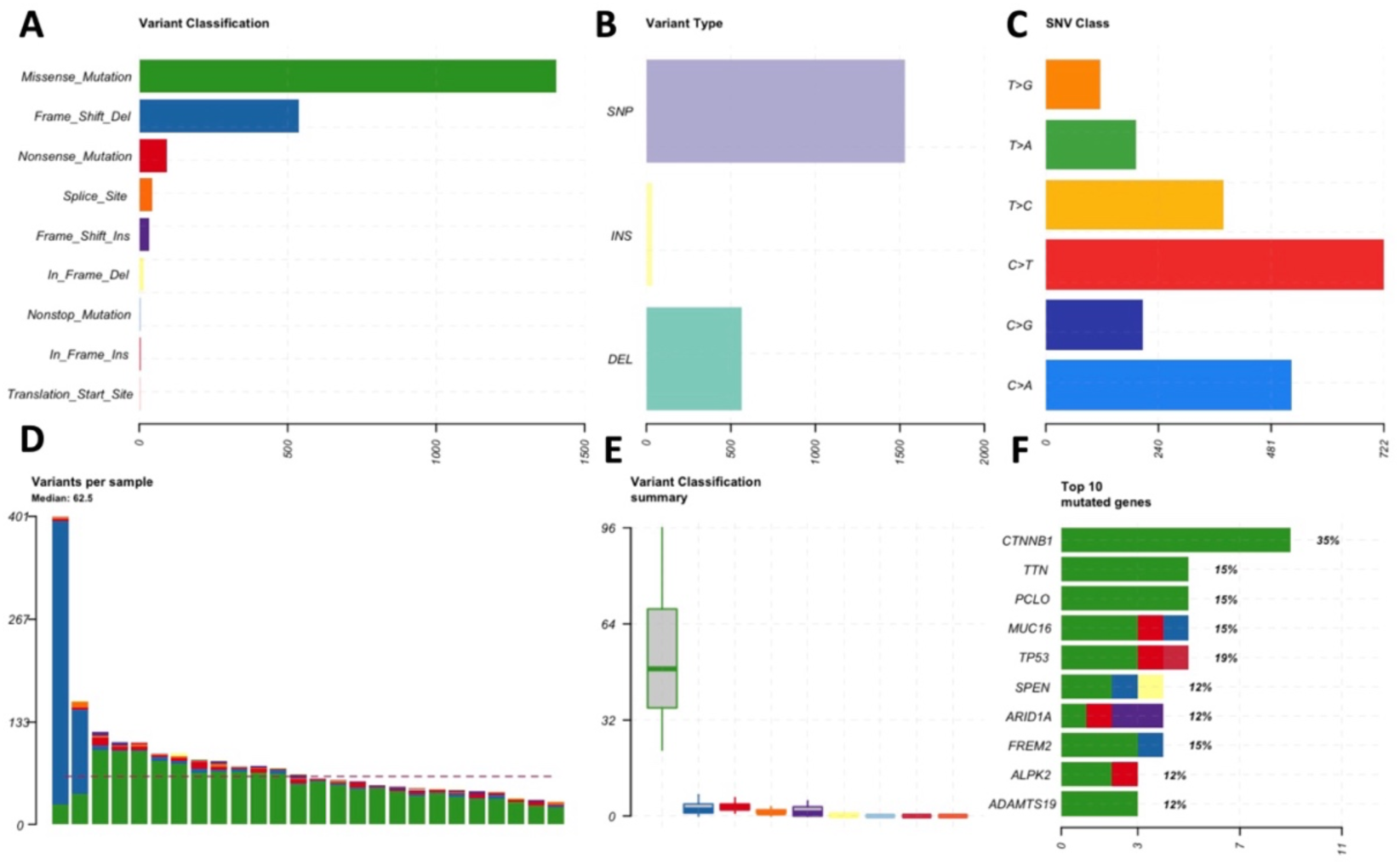
Summary of mutational characteristics of male samples. It includes barplots accounting for number of variants by class (A), type (B) and single nucleotide variation class (C) found in the samples. D) Number of variants per sample (from most to least mutated). E) Boxplots of the number of variants by class, without outliers. F) Top 10 mutated genes, ordered by number of mutations, with percentage of samples in which they are mutated.

#### MIRROR2 phase 1

The miRNA differential network (Fig. 8) has a density of 0.016, where 49% of the edges are negative (representing correlation that were stronger in women) and 51% are positive (correlation that were stronger in men). The 5% most connected miRNAs are: miR-23a-3p, miR-27a-3p, miR-30a-5p, miR-30a-3p, miR-100-5p, miR-181c-5p, miR-144-3p, miR-92b-3p, miR-30c-2-3p, miR-127-5p and miR-654-3p. By gathering the first neighbors of these 11 miRNAs we were able to identify 96 miRNAs of interest to serve as input for the second phase of the algorithm.

**Figure 8.**
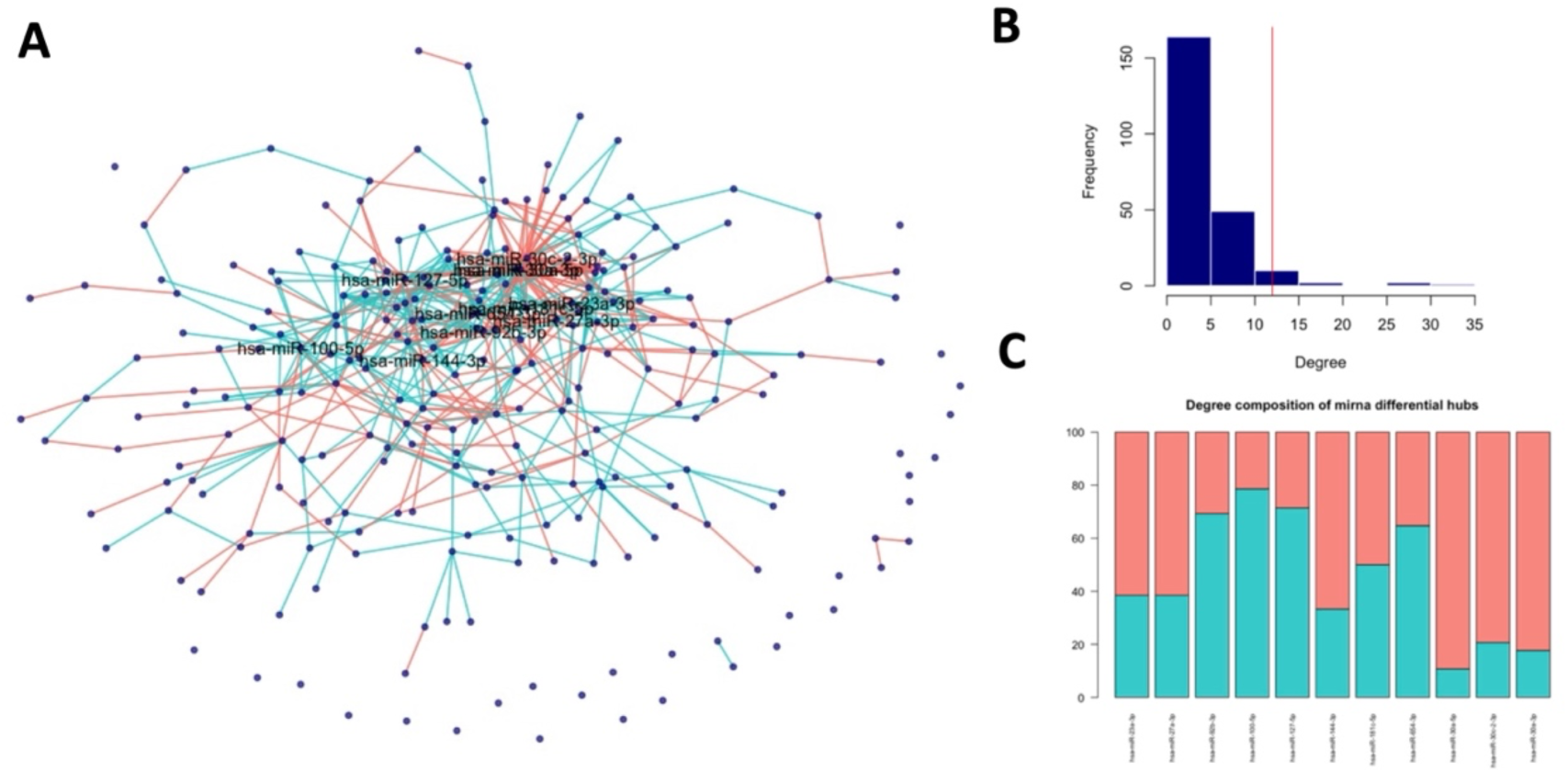
A) miRNA differential co-expression network among males and females. Each node is a miRNA, and each edge is a significant change in co-expression. Blue edges code for a positive differential co-expression (meaning that co-expression is stronger in males) while pink edges code for a negative differential co-expression (meaning that co-expression is stronger in females). B) Degree distribution of network’s nodes. The red vertical line signals the 95th percentile of the distribution. C) Degree composition of the most connected nodes, showed as percentage over total degree.

#### MIRROR2 phase 2

A total of 427 target genes were enriched in the miRNAs identified by Phase1, and 381 of them were available in our mRNA dataset. These target genes are enriched in KEGG pathways such as “Proteoglycans in cancer” (adj-pval = 5.566 · 10^-7^), “Central carbon metabolism in cancer” (adj-pval = 1.45 · 10^-5^), “Wnt signaling pathway” (adj-pval = 5.62 · 10^-4^), “Estrogen signaling pathway” (adj-pval = 1.59 · 10^-3^), highlighting their role in estrogen-mediated processes as well as in metabolic pathways which show sexually dimorphic behaviors. Moreover, these target genes are also enriched in the “FoxO signaling pathway” (adj-pval = 3.30 · 10^-8^ ) which have seen, by experiments on mice, to control the estrogen-dependent resistance and androgen-mediated facilitation of HCC, making it a central process in the sexual dimorphism of HCC^7^.

When inferring differential co-expression networks of case and control samples for males and females separately, we obtained two networks with slightly different density (0.04 for males and 0.02 for females), but a similar degree distribution. Again, the two sets of nodes with the higher strength for the male and female networks only have one common element (Fig. 9A). Moreover, the female network has a balanced amount of positive and negative edges, while the male network has a predominance of negative edges (61%), representing the biggest number of changes are those of correlations that are becoming weaker from control cases to cancer ones. The 19 genes with higher strength in the female differential network are: FANCL, PNMA1, ASB3, ZEB2, BMT2, MBNL2, TIAM1, EZH2, CNOT9, RANBP3, BTBD1, IGF1R, NUFIP2, PLEKHO2, SKIDA1, RBM27, SKP2, SH3GL1, UTP4.

**Figure 9.**
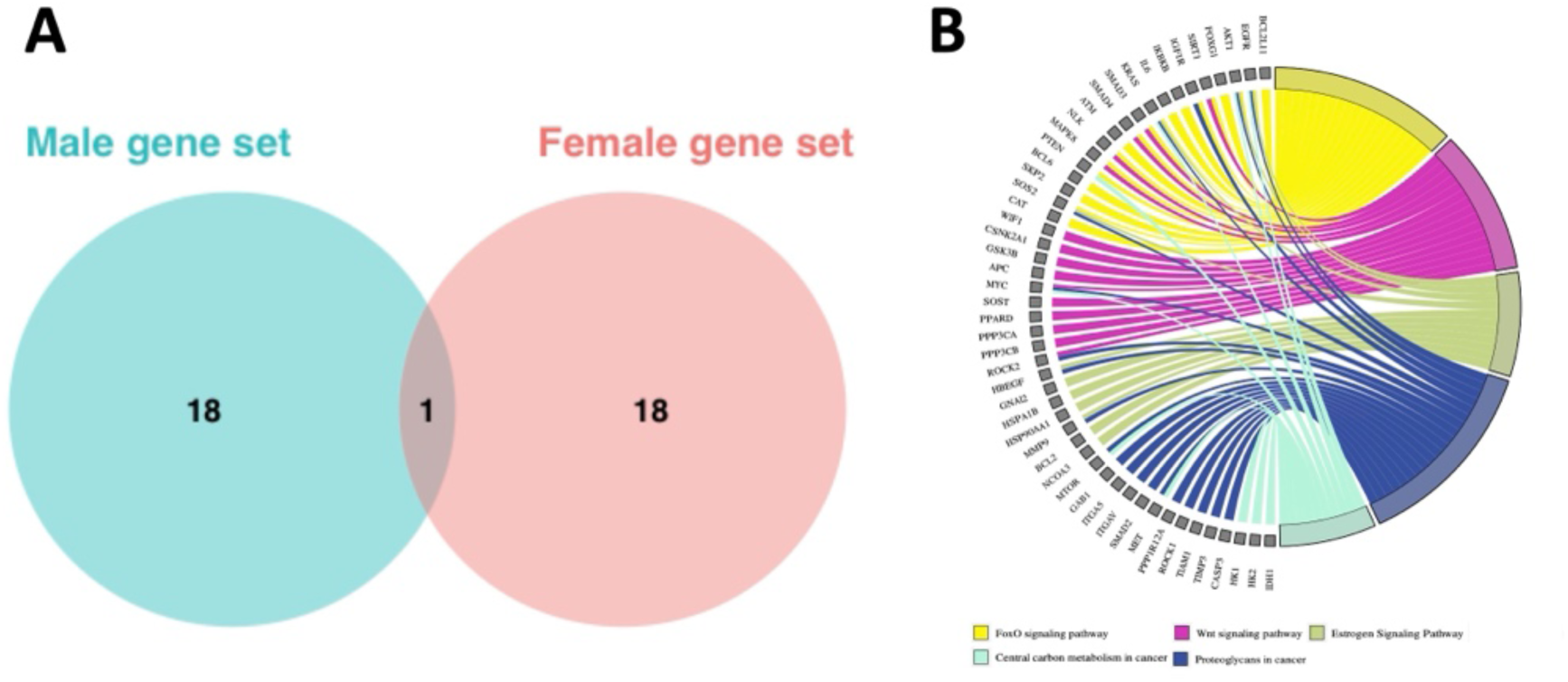
A) Venn diagram showing the overlap between the most connected nodes of the female and male differential co-expression network. B) Chord plot showing the genes belonging to KEGG pathway of interest for this application

For the male differential network instead, they are: SPRY2, SLC38A2, FAM8A1, DYRK1A, NDE1, JDP2, MTDH, INCENP, WDR82, BCL6, GJA1, FNIP1, MIA3, GM2A, CHD1, CNEP1R1, SKP2, GNAL, PPP1R15B.

### Low-grade gliomas

The low-grade gliomas input dataset is made of 310 miRNAs and 128 cancer patients (70 males and 58 females). Patients do not show significant differences in terms of clinical characteristics or survival (Supplementary Figure 3) based on their sex. A similar mutational profile is shown by analyzing variants and SNV among the two sexes; most top mutated genes are also common (Fig. 10 and 11).

**Figure 10.**
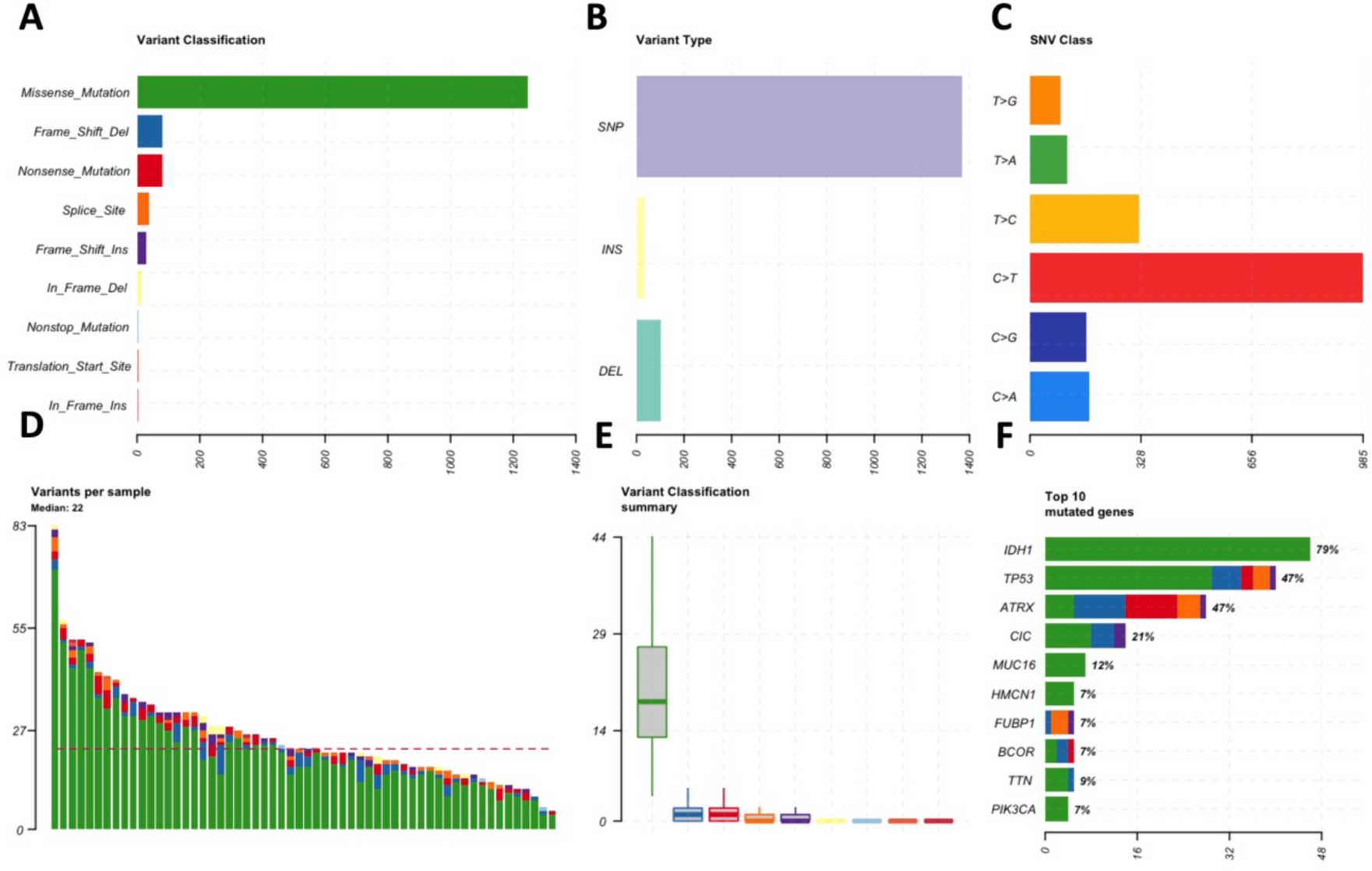
Summary of mutational characteristics of females (above) and male (below) samples. It includes barplots accounting for number of variants by class (A), type (B) and single nucleotide variation class (C) found in the samples. D) Number of variants per sample (from most to least mutated). E) Boxplots of the number of variants by class, without outliers. F) Top 10 mutated genes, ordered by number of mutations, with percentage of samples in which they are mutated.

**Figure 11.**
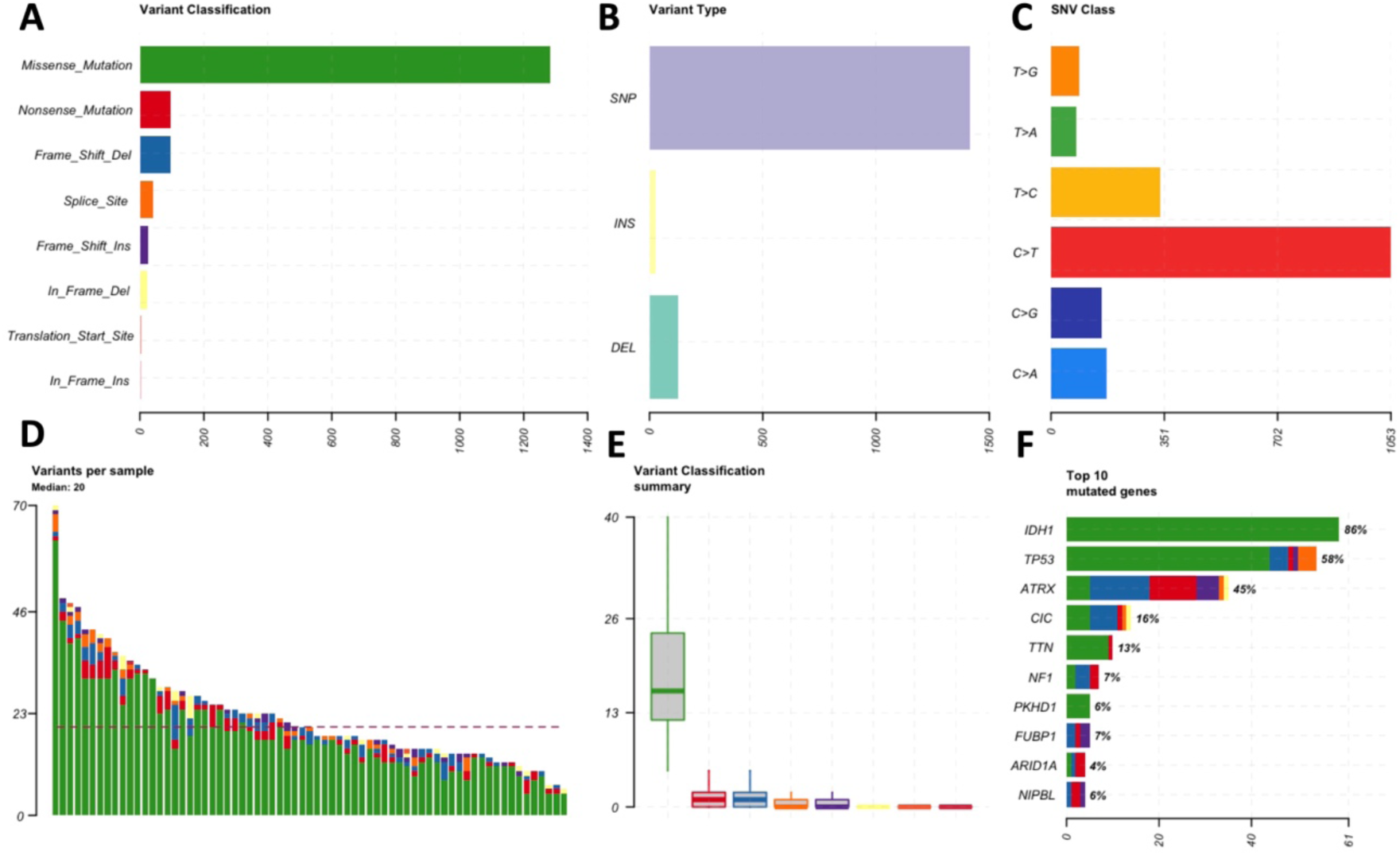
Summary of mutational characteristics of females (above) and male (below) samples. It includes barplots accounting for number of variants by class (A), type (B) and single nucleotide variation class (C) found in the samples. D) Number of variants per sample (from most to least mutated). E) Boxplots of the number of variants by class, without outliers. F) Top 10 mutated genes, ordered by number of mutations, with percentage of samples in which they are mutated.

#### MIRROR2 phase 1

The miRNA differential network (Fig. 12A) has a density of 0.025, where 61% of the edges are negative (representing correlation that were stronger in women) and 39% are positive (correlation that were stronger in men). We identified 11 hub miRNAs (Fig. 12 C): miR-23a-3p, miR-27a-3p, miR-135a-5p, miR-301a-3p, miR-328-3p, miR-337-3p, miR-424-5p, miR-451a, miR-22-5p, miR-29b-2-5p, miR-374a-3p. By gathering the hub miRNAs’ first neighbors in the network we were able to identify 179 miRNAs of interest to serve as input for the second phase of the algorithm.

**Figure 12.**
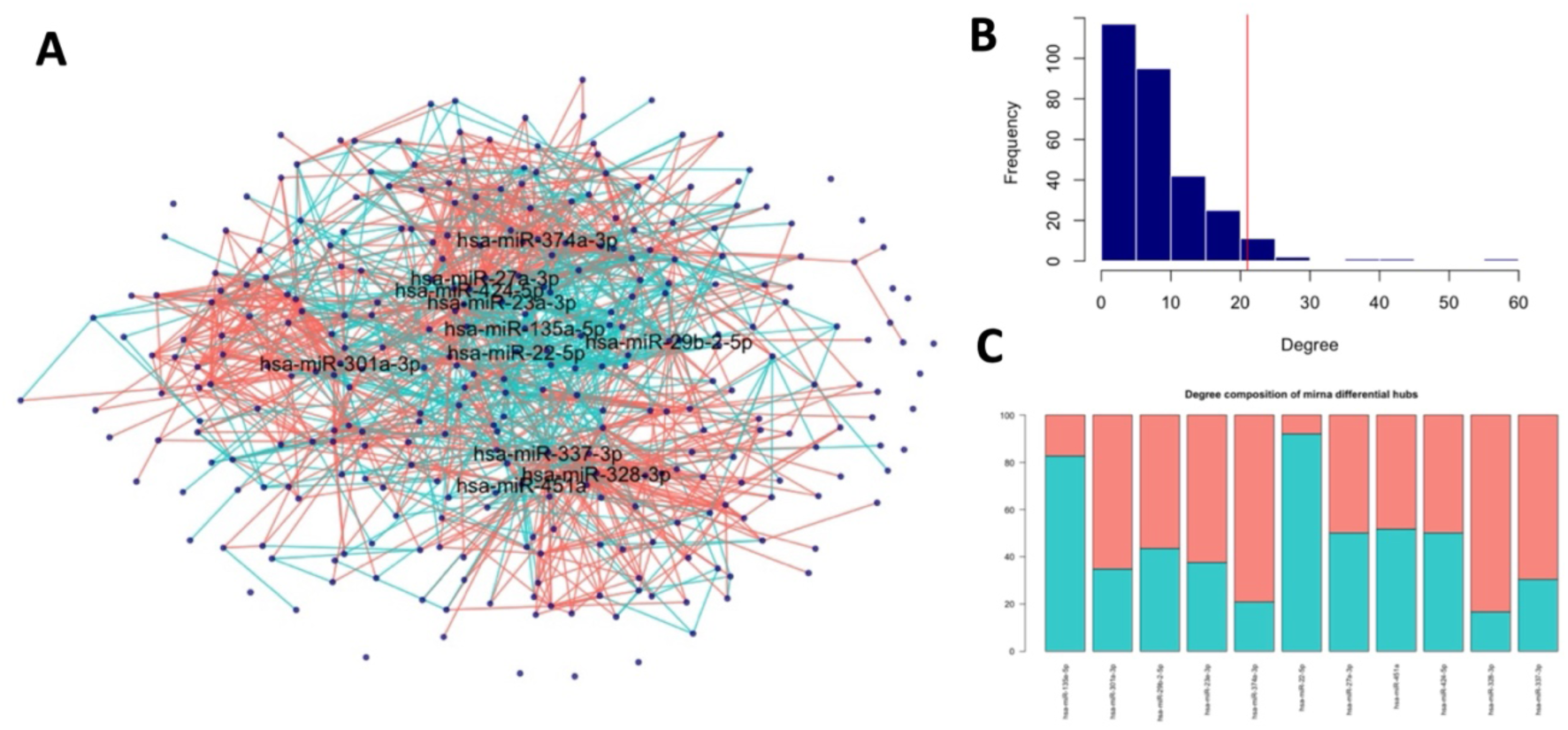
A) miRNA differential co-expression network among males and females. Each node is a miRNA, and each edge is a significant change in co-expression. Blue edges code for a positive differential co-expression (meaning that co-expression is stronger in males) while pink edges code for a negative differential co-expression (meaning that co-expression is stronger in females). B) Degree distribution of network’s nodes. The red vertical line signals the 95th percentile of the distribution. C) Degree composition of the most connected nodes, showed as percentage over total degree

#### MIRROR2 phase 2

A total of 1182 target genes were enriched in the miRNAs identified by Phase1, and 1093 of them were available in our mRNA dataset. Among the results of the KEGG pathways enrichment analysis we found again the “Estrogen signaling pathway” (adj-pval = 2.14 · 10^-4^) as well as the “Lipid and atherosclerosis” pathway (adj-pval = 1.747 · 10^-11^); “multiple preclinical studies described that aberration in lipid metabolism is lipid alteration in glioma, where glioma cells express an increased level of total lipid content compared to normal tissues”, moreover severe dysregulation in phospholipid components has been reported in the IDH mutation subtype^34^. It is also known that the lipid metabolism plays a major role in the regulation of the immune system^35^ and shows sexually dimorphic characteristics^36^. To perform the following steps of the analysis we used control samples from the Gtex Database (see Methods), since paired control samples from the TCGA were not available; after the same preprocessing steps, we obtained 74 healthy female samples and 181 male samples to compare, respectively, to our initial 58 and 70 cancer samples.

When inferring differential co-expression networks of case and control samples for males and females separately, we obtained two networks with density of 0.09 for males and 0.04 for females. Both networks have a balanced amount of positive and negative edges. The output gene sets for the male and female networks have 29 common elements (Fig. 13 A).

**Figure 13.**
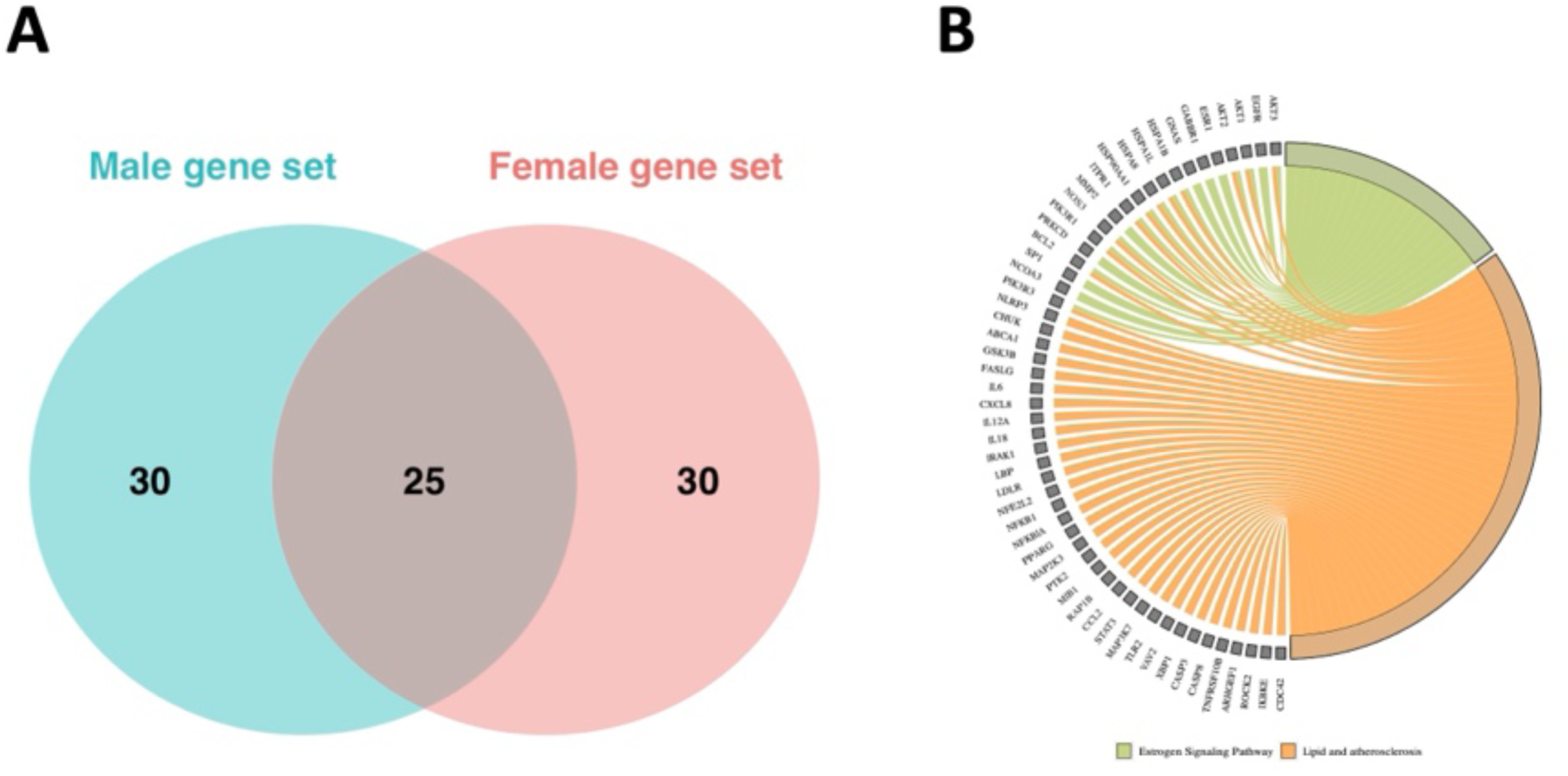
A) Venn diagram showing the overlap between the most connected nodes of the female and male differential co-expression network. B) Chord plot showing the genes belonging to KEGG pathway of interest for this application

The 55 genes with highest strength of the weighted female differential network are: PNPLA4, PLEKHB1, KPNA6, ABCF2, USP28, PLEKHA5, SLK, HADHA, RAPGEF4, MTMR3, ZFYVE21, USP10, BABAM1, RASIP1, GLI3, E2F3, PDGFRB, WNT5A, CLIP4, EPHA4, TGFB3, MXI1, COIL, AMOT, LOXL1, AFDN, HIP1R, MKRN1, PLXNC1, GIT2, SELENBP1, TACC1, SSRP1, MKX, FARP1, GRIA1, EFCAB14, S100B, RAVER1, FYCO1, DLC1, NOS3, FZD6, RET, USP47, SERPINB9, KIF5B, BSG, JUP, PDIK1L, PAWR, TUBB, HDAC2, RNPS1, TUBB3. The 55 genes with highest strength of the weighted male differential network are: PNPLA4, PLEKHB1, KPNA6, ABCF2, PLEKHA5, PITHD1, SLK, HADHA, SIRT1, PSMD8, MTMR3, USP10, CCNE1, BABAM1, KANK1, SMC3, CASC3, EFTUD2, EPHA4, AKT3, MXI1, OCRL, SMARCA4, SH3BP4, CAMSAP1, HIP1R, MYH10, MKRN1, LRP4, PLXNC1, FKBP10, STRIP1, SELENBP1, NOTCH1, MKX, FOXO1, GRIA1, EFCAB14, S100B, NFIA, FYCO1, PRKCD, DLC1, RET, TMEM100, RBPJ, RAB31, KIF5B, PRPF8, PSMD2, YIPF6, SRPRA, DYNC1H1, TUBB3, RASSF5.

In this final application we found a bigger overlap between the output genes for males and females in comparison with the results obtain with first two datasets. We believe this may be a result of not having paired case-control samples; by working with unpaired mRNA data more noise in introduced, reducing MIRROR2’s ability to strongly differentiate between the two sexes.

##### Comparative analysis with other approaches

### Weighted correlation network analysis (WGCNA)

A first validation test was done by applying the WGCNA software; this pipeline revolves around the generation of gene co-expression networks and performing a clustering analysis over them. Module eigengenes (Mes) are then identified as representative nodes of each cluster and used to study the clusters of interest in relation to chosen phenotypes. On the COAD dataset, WGCNA was able to identify two clusters (the “blue cluster” and the “turquoise cluster” of size respectively 70 and 72 genes), while the rest of the nodes remained unassigned (Fig. 14 A). When checking the correlation of these modules with the trait of interest (the sex of the patients, we only found weak correlations (Fig. 14 B). No meaningful module can be detected by the WGCNA; in fact, even by lowering the threshold on the correlation strength and analyzing the obtained modules in terms of module membership (MM) and gene significance (GS) we did not find a strong enough correlation value for both the blue module and the turquoise module (respectively 0.63 and 0.53, Fig. 14 G and H). Finally, in both the HCC and LGG dataset, the WGCNA couldn’t identify meaningful clusters in the networks. In both cases only one cluster (size 53 in HCC and size 75 in LGG) was found and most nodes remained unassigned. Nevertheless, we tried to inspect the correlation of these module the clinical trait of interest (sex), but no meaningful correlation was found.

**Figure 14.**
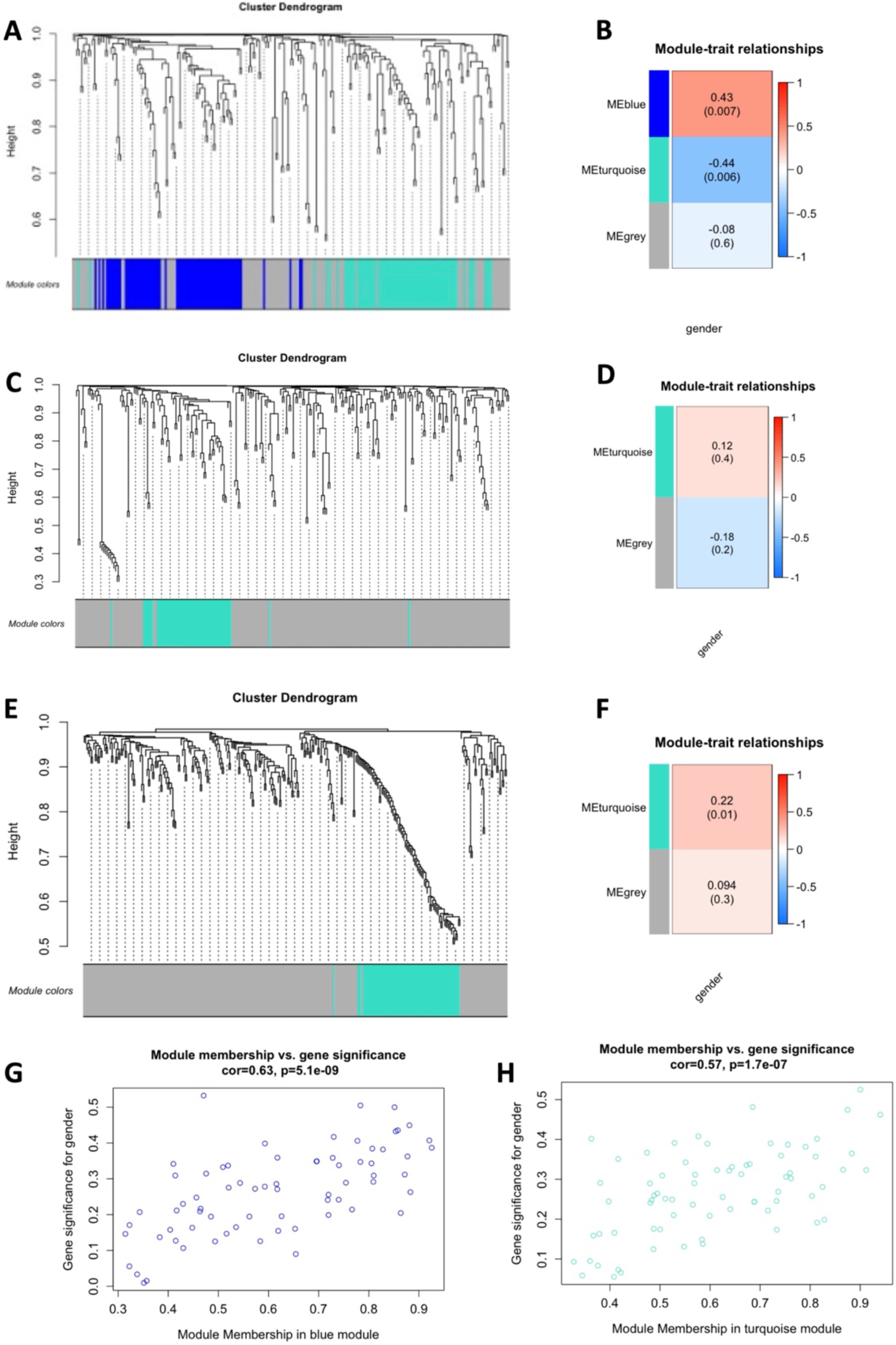
Plots produced by the WGCNA. A, B, G and H refer to the COAD dataset; C and D to LIHC, E and F to LGG

### Differentially expressed genes (DEGs) and miRNAs

#### miRNAs

As a second validation analysis we decided to compare our selection of miRNAs at the end of Phase1 to the one that would have been obtained by performing a differential expression analysis between males and female samples. No differentially expressed miRNAs among the two sexes were found in any of the three datasets used in this work; to make sure the result was not dependent on our choice of threshold we tried softening it till a fold change value of 1. However no significant results were found, proving that this approach would have been successful in this scenario.

#### mRNAs

An additional validation step was carried out by looking for differentially expressed genes among the targets of Phase2. We computed differential expression between cancer and control tissues for women and men separately. For the COAD dataset we found 5 DEGs in men and 15 in women; for the HCC dataset we identified 8 DEGs in men and 7 in women. Finally, for the LGG dataset we found 138 male DEGs and 101 female DEGs. We believe the biggest amount of DEGs in LGG to be related to the use of unpaired samples. Comparing male and females DEGs for each dataset, we saw that they have a big overlap. In fact, by inspecting the correlation of the gene’s fold changes among males and females, we found them to be highly correlated in all three datasets (rho = 0.9 in COAD, rho = 0.78 in HCC, rho = 0.97 in LGG) (Fig. 15– A, B, C).

**Figure 15.**
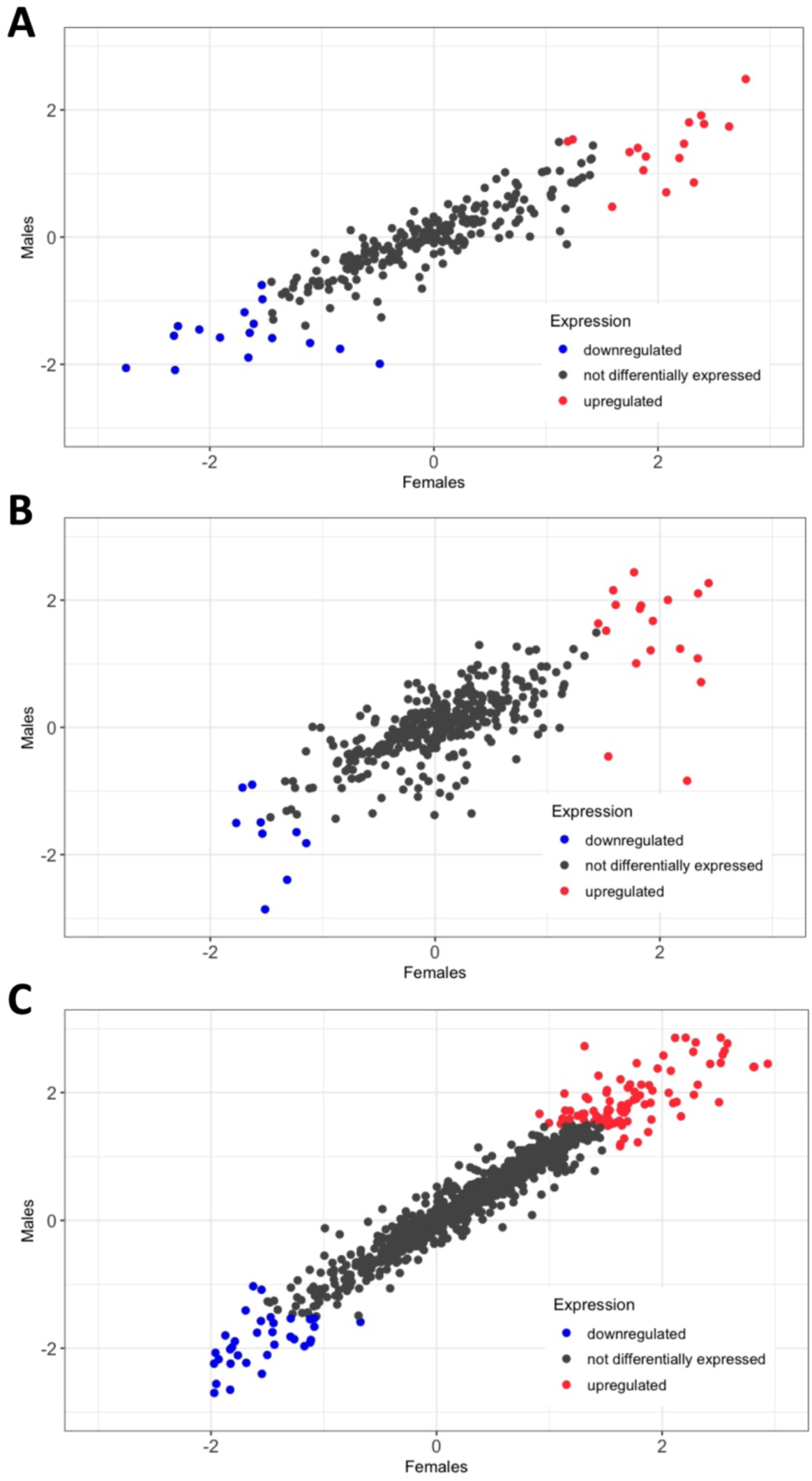
Scatterplots showing the correlation of gene fold changes computed in males and females. Genes which are upregulated in at least one of the two sets are red; genes which are downregulated in at least one of the two sets are blue (A for COAD, B for HCC, C for LGG)

#### Evaluation of robustness with respect to the sex variable

To assess how specific our miRNAs of interest are to the sex-based differential co-expression network, we performed permutations on the input dataset to obtain 1000 random networks, from which we retrieved the most connected miRNAs and their first neighbors. We saw that the median overlap coefficient among our set of miRNAs and the ones obtained from the random networks is 0 is all three datasets, while the overlap with the simulated sets of first neighbors is significative in less the 10% of the cases (2.5% in COAD, 9.5% in LIHC and 1.8% in LGG)

### Clinical validation

To show the relevance of the genes identified by MIRROR2, we built logistic regression models to predict the survival outcome (high or low) of a patient, using the expression of these genes in the cancer samples as predictors. This was done separately for males and females of the TGCA-COAD dataset, using the sex-specific genes given as output by MIRROR2. The female model relies on 12 genes, and it was trained on 166 patients and tested on 41 (20% of all female patients). It has a ROC/AUC score of 0.68 and an F1 score of 0.66 (Fig. 16) The male model also relies on 12 genes; it was trained on 187 patients and tested on 46. It has a ROC/AUC score of 0.62 and an F1 score of 0.66. Moreover, to make sure that our models, and the genes they based on, were sex-specific we tried applying the female model on male data and vice versa. Doing so, we proved that the female model does not perform well on male data (ROC/AUC = 0.57, F1= 0.41). Similarly, the male model is not as good as the female model on female data (ROC/AUC = 0.57, F1 = 0.55).

**Figure 16.**
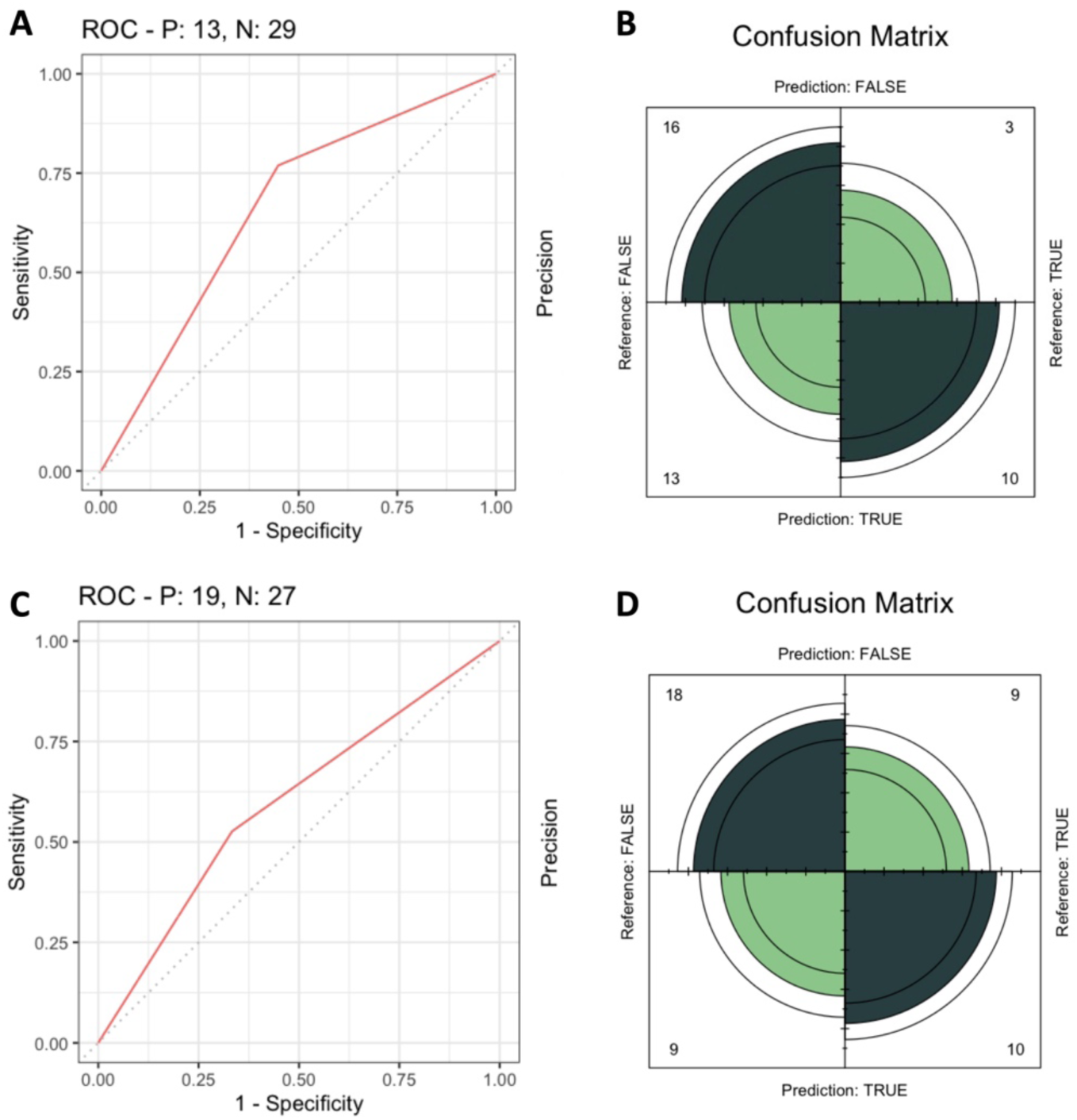
ROC curve and confusion matrix of the logistic regression models trained on female data (A, B) and male data (C, D) to predict survival expectancy (higher or lower than 2 years)

A final validation was done by selecting a set of 12 random genes from our mRNA dataset (excluded the ones identified by MIRROR2) for 1000 times and using the to train and test models. On average the performance obtained by these models in terms of F1 score is around 0.6, lower than that obtained by using MIRROR2’s output genes.

These results highlight the predictive power held by the genes identified my MIRROR2. By choosing alternate classification algorithm, performing fine tuning and possibly using a bigger dataset we expect that the accuracy would increase.

## Discussion and conclusions

Having already established the importance of sex-specific medicine in shaping health-care solutions that are optimal for both males and females, especially in oncology, it becomes clear that there is an urgent need to explore the complex molecular mechanisms that are behind these disparities. Unique effects that can be traced back to a patient’s sex can be seen on both cancer development and progression, as well as in treatment response. The evolving landscape of precision oncology has already provided compelling evidence of sex disparities in cancer that extend far beyond reproductive tissues; however, comprehensively understanding the molecular mechanisms that drive sex-specific characteristics of cancer remains an open challenge. In this context, defining a strategy to investigate these differences and identifying sex-specific molecules to focus research efforts on presents a promising opportunity for identifying novel biomarkers, prognostic indicators, and/or therapeutic targets tailored to individual patients. For this purpose, we proposed MIRROR2 (an updated version of our algorithm MIRROR^26^), which integrates expression data of both miRNAs and mRNAs to study sex differences in cancer molecular mechanisms at two different regulatory levels with the goal of identifying sex-specific key genes. MIRROR2 is not only able to identify molecules with a sex-biased behavior thanks to the integration of two molecular layers, but it goes beyond a single-molecule approach; indeed, it focuses of networks of interrelationships, benefiting from a more comprehensive and holistic view of the biological processes at play. By analyzing sets of miRNAs and/or mRNAs at the same time and how their co-expressions patterns differ, the final output of MIRROR2 is the result of a multi-layer analysis of the complex molecular phenomena that characterize cancer. In fact, while the first version of MIRROR ended after the miRNAs’ targets retrieval and their functional analysis, MIRROR2 uses differential co-expression analysis to explore a second molecular layer: mRNA expression. By comparing cancer and control tissues in both women and men separately, MIRROR2 is able to identify molecules which have a key and sex-specific role in cancer biology. Moreover, MIRROR2 has a more refined way of selecting the key miRNAs from the M-vs-F-differential co-expression network which can identify a more compact set of miRNAs showing a strong sexually dimorphic behavior.

To assess the performance of our algorithm we tested it on three datasets related to three different tumors: colon adenocarcinoma, hepatocellular carcinoma and low-grade gliomas. On the same datasets we also applied other well-known methods commonly used to identify key genes.

Across all three datasets, MIRROR2 has been able to identify sex-specific key genes (one set for female patients and another one for male patients) which are involved in numerous co-expression changes between case and control samples and are part of a wider network of genes involved in sex-biased pathways. Moreover, the two sets of genes identified in each dataset are almost completely different, which means that they are extremely specific to each sex. On the contrary, applying state of the art methods did not produce the same effect; in fact, by comparing miRNA expression among males and females, no differentially expressed miRNAs were found in all three datasets. This highlights the importance of going beyond measuring expression variation as a way to compare groups of samples. According to this approach no miRNA-related difference would have been found in these three datasets. However, after applying MIRROR2 we know that, even if differential expression among males and females is not significantly different, the co-expression interrelationships of the miRNA differ in many ways. The WGCNA was also unfit for the purposes of this analysis; in fact, when using it to compare weighted co-expression modules of miRNAs, it failed to identify a good community structure of the networks, resulting in most nodes not being assigned to any cluster and, in two out of three cases, to just one cluster being found. Moreover, the identified clusters showed only weak or nonexistent correlation to the patients’ sex, failing to suggest strong key genes candidates.

Even when testing out the computation of differentially expressed genes after having performed Phase1 of our algorithm, we saw that males and females preset a very similar (and in some cases almost identical) profile of differential gene expression. This means that this type of measure would not be useful to identify key genes that are sex-specific, even when starting out from a set of mRNAs associated to the miRNAs previously selected by MIRROR2. Moreover, to show the clinical relevance of our output genes, we used MIRROR2’s output from the COAD application to build logistic regression models for the prediction of the patient’s survival expectancy (higher or lower than 2 years). We showed not only how these genes have a predictive power over survival and can be integrated with clinical features, but also how this predictive power is sex specific: the model built on the female gene set performed worse on males and vice versa.

In conclusion, we can confidently say that MIRROR2 is a viable option for the identification of sex-specific key genes in oncology and for the study of the complex phenomena involved in sexual dimorphic characteristics of cancer. MIRROR2 is not only able to provide good results, but it is also filling a methodological gap: to our knowledge there is no existing algorithm tailored made for the study of sexual dimorphism, and the current most used approaches on similar topics did not prove successful in this application. Indeed, comparison with other methods showed that pure differential analysis (first-order statistics) is not able to identify differential genes, and that without a specific analysis design involving sex information, these genes cannot be recovered either.

Nevertheless, we have to acknowledge certain limitations; first of all, even though MIRROR2 can be easily scaled-down to bigger input datasets, the fact that it relies on gender-balanced miRNA samples as well as case and control mRNA samples often leads to working with a reduced input size. Moreover, in the applications we tested we worked with data gathered from the TCGA database, which were lacking proper clinical and pathological information on the patients; this prevented us from further exploring the link between MIRROR2’s output genes and clinical data as well as having a more complete understanding of our patients’ condition. For example, it was not possible to check if the considered patients had any kind chromosomal aneuploidies. However, we were able to check that the genes produced as output by MIRROR2 in the three considered cancers are not located in X or Y chromosomes. Further improvements of this method could focus in extending the algorithm testing to more cancer as well as to investing more extensively the connection between the output genes and clinical characteristics.

## Supplementary Figures

**Figure S1.**
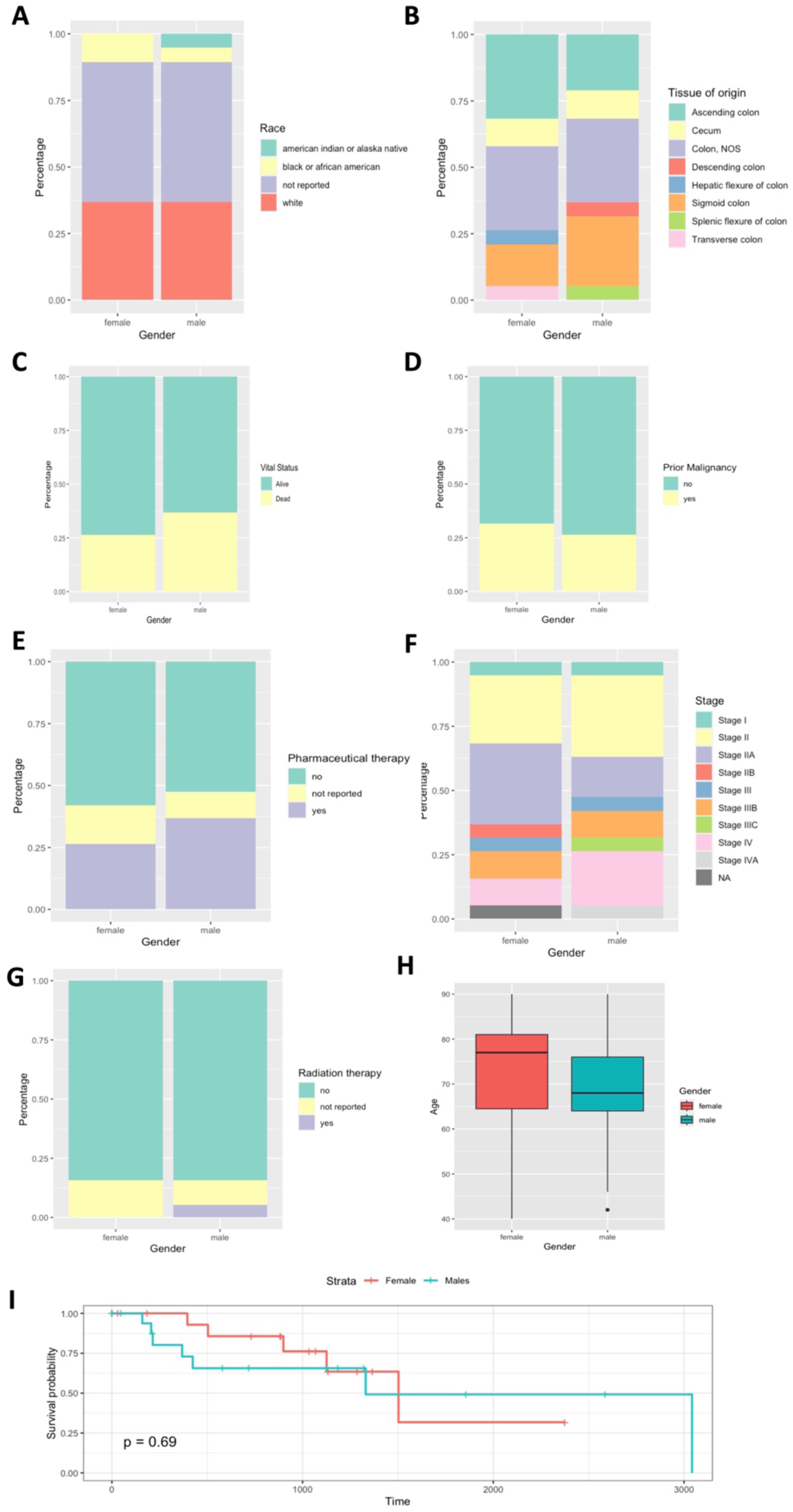
Summary of clinical data of considered patients from the COAD dataset

**Figure S2.**
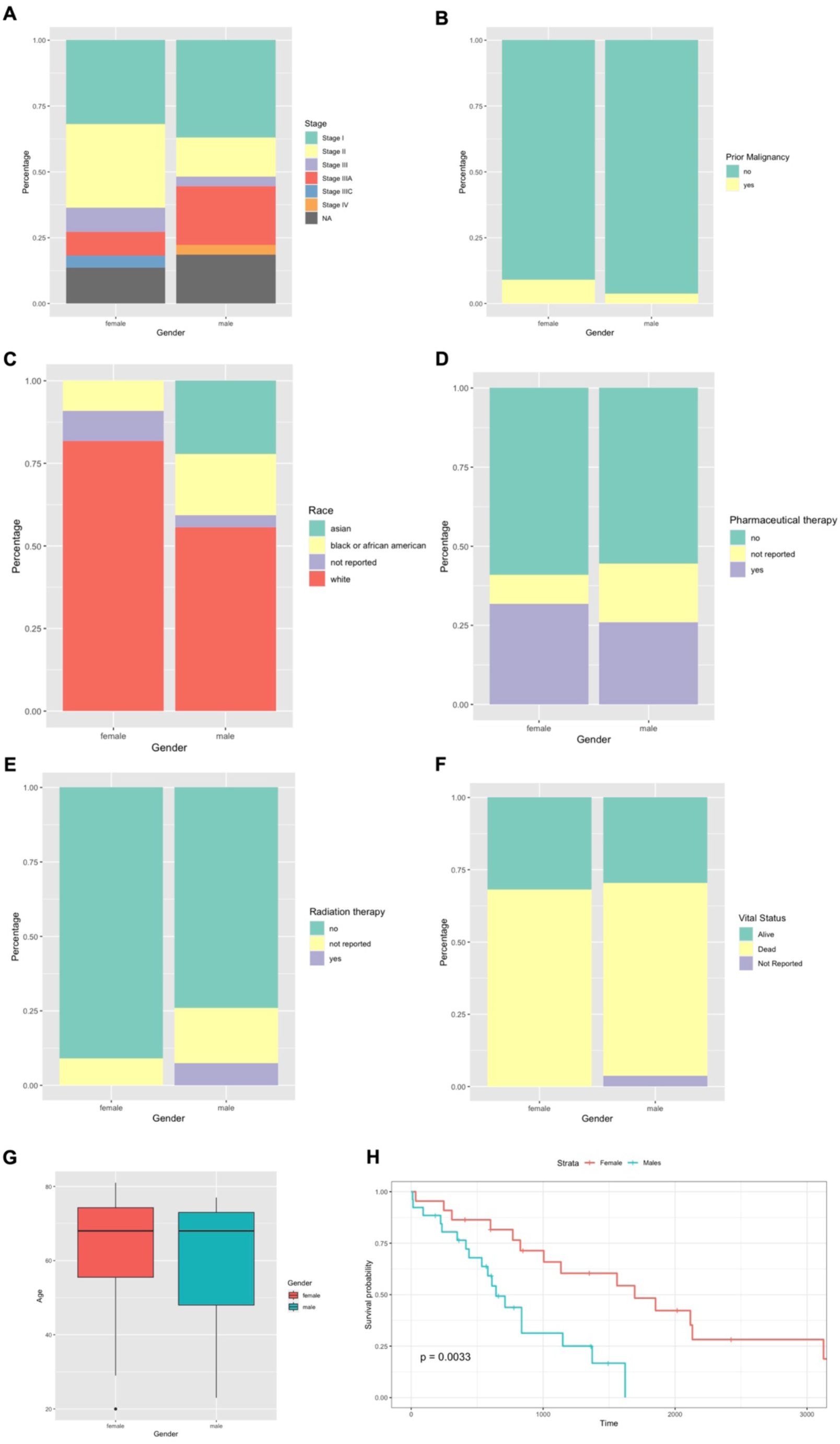
Summary of clinical data of considered patients from the LIHC dataset

**Figure S3.**
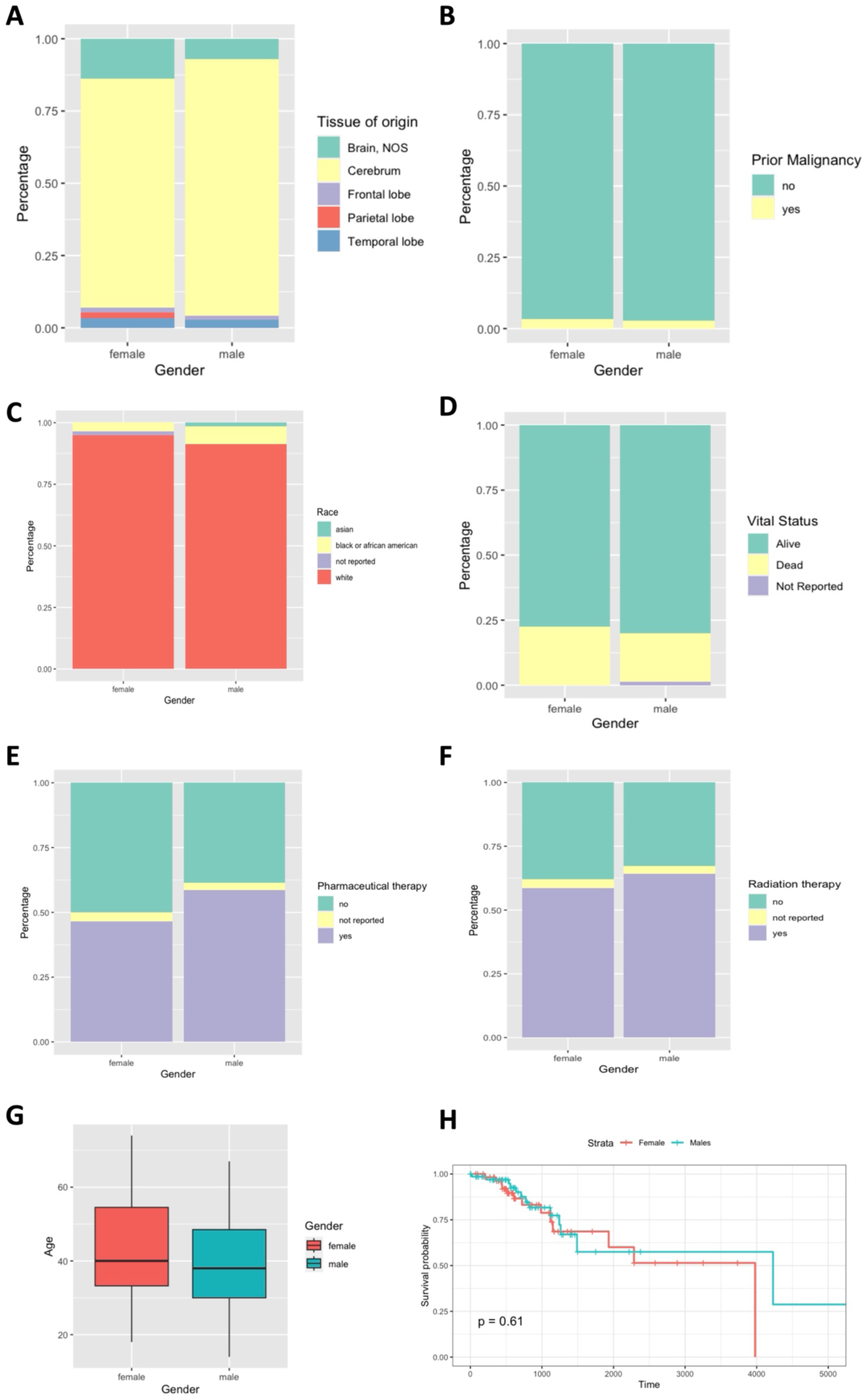
Summary of clinical data of considered patients from the LGG dataset

## Acknowledgements

A. is a fellow of the Phd Network Oncology and Precision Medicine, Department of Experimental Medicine, Sapienza University of Rome.

## Author contributions: CRediT

**Caterina Alfano:** Conceptualization, Methodology, Software, Data Curation, Validation, Investigation, Writing - Original Draft, Visualization

**Marco Filetti:** Writing - Review & Editing, Conceptualization

**Lorenzo Farina:** Supervision, Conceptualization, Writing - Review & Editing

**Manuela Petti:** Conceptualization, Methodology, Investigation, Writing - Review & Editing, Supervision, Project administration, Funding acquisition

## References

1. Wagner AD, Oertelt-Prigione S, Adjei A, et al. Gender medicine and oncology: report and consensus of an ESMO workshop. Ann Oncol. 2019;30(12):1914–1924. doi:10.1093/annonc/mdz414

2. Wolrd health organization - gender and medicine. https://www.who.int/health-topics/gender#tab=tab_1

3. Kim N. Application of sex/gender-specific medicine in healthcare. Korean J Women Health Nurs. 2023;29(1):5–11. doi:10.4069/kjwhn.2023.03.13

4. Dart A. Sexual dimorphism in cancer. Nat Rev Cancer. 2020;20(11):627–627. doi:10.1038/s41568-020-00304-2

5. White A, Ironmonger L, Steele RJC, Ormiston-Smith N, Crawford C, Seims A. A review of sex-related differences in colorectal cancer incidence, screening uptake, routes to diagnosis, cancer stage and survival in the UK. BMC Cancer. 2018;18(1):906. doi:10.1186/s12885-018-4786-7

6. Barzi A, Lenz AM, Labonte MJ, Lenz HJ. Molecular Pathways: Estrogen Pathway in Colorectal Cancer. Clin Cancer Res. 2013;19(21):5842–5848. doi:10.1158/1078-0432.CCR-13-0325

7. Li Z, Tuteja G, Schug J, Kaestner KH. Foxa1 and Foxa2 Are Essential for Sexual Dimorphism in Liver Cancer. Cell. 2012;148(1-2):72–83. doi:10.1016/j.cell.2011.11.026

8. Klein SL, Flanagan KL. Sex differences in immune responses. Nat Rev Immunol. 2016;16(10):626–638. doi:10.1038/nri.2016.90

9. Beery AK, Zucker I. Sex bias in neuroscience and biomedical research. Neurosci Biobehav Rev. 2011;35(3):565–572. doi:10.1016/j.neubiorev.2010.07.002

10. Shireman JM, Ammanuel S, Eickhoff JC, Dey M. Sexual dimorphism of the immune system predicts clinical outcomes in glioblastoma immunotherapy: A systematic review and meta-analysis. Neuro-Oncol Adv. 2022;4(1):vdac082. doi:10.1093/noajnl/vdac082

11. Yasinjan F, Xing Y, Geng H, et al. Immunotherapy: a promising approach for glioma treatment. Front Immunol. 2023;14:1255611. doi:10.3389/fimmu.2023.1255611

12. Garufi C, Giacomini E, Torsello A, et al. Gender effects of single nucleotide polymorphisms and miRNAs targeting clock-genes in metastatic colorectal cancer patients (mCRC). Sci Rep. 2016;6(1):34006. doi:10.1038/srep34006

13. Cai Y, Rattray NJW, Zhang Q, et al. Sex Differences in Colon Cancer Metabolism Reveal A Novel Subphenotype. Sci Rep. 2020;10(1):4905. doi:10.1038/s41598-020-61851-0

14. Haupt S, Haupt Y. Cancer and Tumour Suppressor p53 Encounters at the Juncture of Sex Disparity. Front Genet. 2021;12:632719. doi:10.3389/fgene.2021.632719

15. Guo L, Zhang Q, Ma X, Wang J, Liang T. miRNA and mRNA expression analysis reveals potential sex-biased miRNA expression. Sci Rep. 2017;7(1):39812. doi:10.1038/srep39812

16. A Eaves L, Phookphan P, E Rager J, et al. A Role for microRNAs in the Epigenetic Control of Sexually Dimorphic Gene Expression in the Human Placenta. Epigenomics. 2020;12(17):1543–1558. doi:10.2217/epi-2020-0062

17. Ahmed A, Dai R. Sexual dimorphism of miRNA expression: a new perspective in understanding the sex bias of autoimmune diseases. Ther Clin Risk Manag. Published online March 2014:151. doi:10.2147/TCRM.S33517

18. Lee YS, Dutta A. MicroRNAs in Cancer. Annu Rev Pathol Mech Dis. 2009;4(1):199–227. doi:10.1146/annurev.pathol.4.110807.092222

19. Hasáková K, Bezakova J, Vician M, Reis R, Zeman M, Herichova I. Gender-Dependent Expression of Leading and Passenger Strand of miR-21 and miR-16 in Human Colorectal Cancer and Adjacent Colonic Tissues. Physiol Res. Published online December 29, 2017:S575–S582. doi:10.33549/physiolres.933808

20. Lopes-Ramos CM, Chen CY, Kuijjer ML, et al. Sex Differences in Gene Expression and Regulatory Networks across 29 Human Tissues. Cell Rep. 2020;31(12):107795. doi:10.1016/j.celrep.2020.107795

21. Bai P song, Hou P, Kong Y. Hepatitis B virus promotes proliferation and metastasis in male Chinese hepatocellular carcinoma patients through the LEF-1/miR-371a-5p/SRCIN1/pleiotrophin/Slug pathway. Exp Cell Res. 2018;370(1):174–188. doi:10.1016/j.yexcr.2018.06.020

22. Li C, Yeh K, Liu W, et al. Elevated p53 promotes the processing of miR-18a to decrease estrogen receptor-α in female hepatocellular carcinoma. Int J Cancer. 2015;136(4):761–770. doi:10.1002/ijc.29052

23. Kwekel JC, Vijay V, Han T, Moland CL, Desai VG, Fuscoe JC. Sex and age differences in the expression of liver microRNAs during the life span of F344 rats. Biol Sex Differ. 2017;8(1):6. doi:10.1186/s13293-017-0127-9

24. Hong ES, Wang SZ, Ponti AK, et al. miR-644a is a tumor cell-intrinsic mediator of sex bias in glioblastoma. Published online March 12, 2024. doi:10.1101/2024.03.11.584443

25. Velázquez-Vázquez D, Del Moral-Morales A, Cruz-Burgos J, Martínez-Martínez E, Rodríguez-Dorantes M, Camacho-arroyo I. Expression analysis of progesterone-regulated miRNAs in cells derived from human glioblastoma. Mol Med Rep. 2021;23(6):475. doi:10.3892/mmr.2021.12114

26. Alfano C, Filetti M, Farina L, Petti M. MIRROR: miRNA regulation-level differential network to study sex and ethnic disparities in cancer. In: 2024 46th Annual International Conference of the IEEE Engineering in Medicine and Biology Society (EMBC). IEEE; 2024:1–4. doi:10.1109/EMBC53108.2024.10782801

27. Xu T, Su N, Liu L, et al. miRBaseConverter: an R/Bioconductor package for converting and retrieving miRNA name, accession, sequence and family information in different versions of miRBase. BMC Bioinformatics. 2018;19(S19):514. doi:10.1186/s12859-018-2531-5

28. Michael Love SA. DESeq2. Published online 2017. doi:10.18129/B9.BIOC.DESEQ2

29. Fukushima A, Nishida K. DiffCorr: Analyzing and Visualizing Differential Correlation Networks in Biological Data. Published online April 2, 2015:0.4.4. doi:10.32614/CRAN.package.DiffCorr

30. Langfelder P, Horvath S. WGCNA: an R package for weighted correlation network analysis. BMC Bioinformatics. 2008;9(1):559. doi:10.1186/1471-2105-9-559

31. Fernández-Ponce C, Geribaldi-Doldán N, Sánchez-Gomar I, et al. The Role of Glycosyltransferases in Colorectal Cancer. Int J Mol Sci. 2021;22(11):5822. doi:10.3390/ijms22115822

32. Hartman J, Edvardsson K, Lindberg K, et al. Tumor Repressive Functions of Estrogen Receptor β in SW480 Colon Cancer Cells. Cancer Res. 2009;69(15):6100–6106. doi:10.1158/0008-5472.CAN-09-0506

33. Abancens M, Bustos V, Harvey H, McBryan J, Harvey BJ. Sexual Dimorphism in Colon Cancer. Front Oncol. 2020;10:607909. doi:10.3389/fonc.2020.607909

34. Abdul Rashid K, Ibrahim K, Wong JHD, Mohd Ramli N. Lipid Alterations in Glioma: A Systematic Review. Metabolites. 2022;12(12):1280. doi:10.3390/metabo12121280

35. Garcia C, Andersen CJ, Blesso CN. The Role of Lipids in the Regulation of Immune Responses. Nutrients. 2023;15(18):3899. doi:10.3390/nu15183899

36. Mittendorfer B. Sexual Dimorphism in Human Lipid Metabolism. J Nutr. 2005;135(4):681–686. doi:10.1093/jn/135.4.681

